# Engineering the auxin-inducible degron system for tunable *in vivo* control of organismal physiology

**DOI:** 10.1101/2024.09.05.611487

**Authors:** Jeremy Vicencio, Daisuke Chihara, Matthias Eder, Julie Ahringer, Nicholas Stroustrup

## Abstract

The physiological mechanisms governing health and disease exhibit complex interactions between multiple genes and gene products. To study the dynamics of living systems, researchers need experimental methods capable of producing calibrated, quantitative perturbations *in vivo* — perturbations that cannot be obtained using classical genetics, RNAi interference, or small molecule drugs. Recently, an auxin-inducible degron (AID) system has been developed to allow targeted degradation of proteins using small-molecule activators, providing spatiotemporal control of protein abundance. However, a better understanding of the biochemical activities of AID system components in their physiological context is needed to design quantitative interventions.

Here, we apply engineering approaches to characterize and understand the performance of several AID technologies and then improve this performance in the multicellular animal *Caenorhabditis elegans*. We 1) develop new technologies that allow for a careful calibration of AID activity for specific purposes; 2) develop new TIR1 enzyme constructs with improved performance over existing constructs; 3) develop an approach to simultaneously and independently degrade target proteins in distinct tissues; and finally, 4) develop an approach for pan-organismal protein degradation by re-engineering the TIR1 enzyme. Taken together, these advances enable new quantitative experimental approaches to study the cellular and systems dynamics of animals.

## INTRODUCTION

Experimental manipulation of gene activities is the central approach in molecular genetics, allowing researchers to directly establish the causal role of genes in determining phenotypes. Many biological systems exhibit complex quantitative dynamics, and probing these dynamics requires advanced experimental interventions that can subtly perturb system dynamics in a controlled way (Adamson et al., 2016; McIsaac et al., 2013; Otoupal et al., 2017). To this end, a diverse set of biotechnological approaches have been developed to facilitate spatiotemporal control of gene expression, including recombination with the Cre-lox (Sauer & Henderson, 1988) and FLP-FRT (Cherepanov & Wackernagel, 1995) systems, promoter manipulation using CRISPR-Cas genome editing (Frati et al., 2024; Yao et al., 2024) RNA interference (RNAi) (Fire et al., 1998), or protein degradation using small-molecule proteolysis targeting chimeras (PROTACs) (Buckley et al., 2015).

Another approach, auxin-inducible degradation (AID) (Nishimura et al., 2009) has gained rapid adoption due to many practical advantages including rapid and reversible spatiotemporal control. The AID system evolved in plants as a mechanism through which the hormone auxin regulates proteins required for growth and development (Teale et al., 2006). Adapted first into yeast and a wide range of eukaryotic cells (Nishimura et al., 2009), then *C. elegans* (L. Zhang et al., 2015), *Drosophila* (Trost et al., 2016), and mice (Macdonald et al., 2022), researchers have found that two components of the AID system are sufficient to support targeted protein degradation in animals: first, the F-box transport inhibitor response 1 (TIR1) protein which interacts with the Skp1-Cul1-F-box (SCF) E3 ligase complex to polyubiquitylate target proteins, and second, a short amino acid tag (called a “degron”) that is fused to target proteins and acts as a TIR1 substrate (Tan et al., 2007, Fig. 1a). The TIR1 induces ubiquitylation of degron-tagged proteins only in the presence of the small-molecule indole-3-acetic acid (IAA), providing a means for small-molecule-induced proteasomal degradation (Gray et al., 2001). The high degree of conservation of Skp1 and Cul1 among eukaryotes allows transgenic TIR1 to interact with the Skp1 and Cul1 proteins endogenous to the host species (Bloom et al., 2006; Schulman et al., 2000), removing the need for transgenic E3-Ubiquitin ligases.

**Figure 1.**
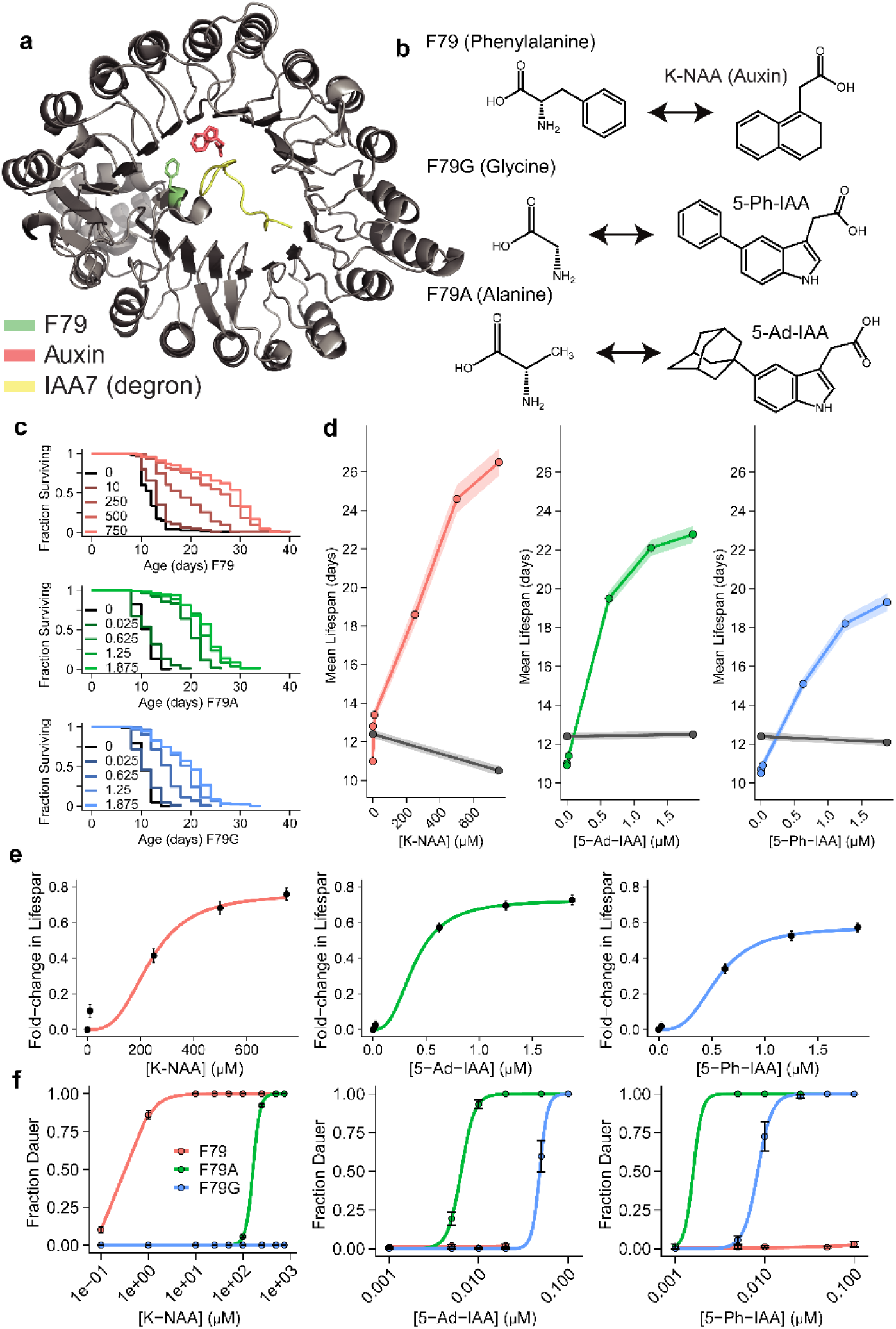
The structure and organismal activities of three TIR1 variants. (a) The crystal structure of AtTIR1 (adapted from Tan et al., 2007). (b) The three amino acids considered in the auxin-binding pocket of TIR1, along with their high-affinity ligands. (c) Kaplan-Meier survival curves showing the dose-response of lifespan of K-NAA with TIR1[F79] (red), 5-Ad-IAA with F79A (green), and 5-Ph-IAA with F79G (blue), in populations expressing DAF-2::AID and the respective TIR1 variant. (d) The mean (points) and standard error (shaded region) of the same populations (colors) compared to wildtype (black). (e) The fold-change in lifespan observed in the same populations, fit by Hill functions. (f) The same TIR1[F79] (red), F79A (green), and F79G (blue) strains but this time, assaying for a constitutive Dauer entry phenotype on K-NAA (left), 5-Ad-IAA (center), and 5-Ph-IAA (right).

The AID system has been shown to be capable of depleting most of the target protein between 15 minutes and 3 hours depending on the half-life of the target protein (Holland et al., 2012; Nishimura et al., 2009). In addition, this depletion is rapidly reversible, with recovery time being inversely proportional to the concentration of auxin used (L. Zhang et al., 2015). In contrast, RNA interference techniques involve a significant time delay between induction and depletion of the target (Neofytou et al., 2017) and is not necessarily reversible due to the relatively long half-life of the small interfering RNAs (siRNAs) once loaded into the RNA-induced silencing complex (Bartlett & Davis, 2006).

A variety of variants of the AID system have been developed to optimize its performance and enable new experimental techniques. In plants, botanists designed an orthogonal small-molecule–TIR1 pair by employing a “bump-and-hole” strategy (Uchida et al., 2018) that alters the TIR1 ligand-binding pocket to recognize a chemically-modified auxin derivative, providing a means for target protein degradation independent from the endogenous AID system. This system was subsequently adapted into animals, branded as the “AID version 2” (AID2) (Yesbolatova et al., 2020). In this system, a single amino acid substitution, F79G, dramatically increases the affinity of *Arabidopsis thaliana* TIR1 (F74G in *Oryza sativa* TIR1) to the synthetic auxin analog: 5-phenyl-indole-3-acetic acid (5-Ph-IAA) (Fig. 1b). TIR1[F79G] ubiquitination activity can be induced at 670-fold lower concentrations of 5-Ph-IAA compared to those required of IAA to activate the original TIR1[F79]. The AID2 system has been reported to have faster depletion kinetics compared to the original system and shows lower “basal” auxin-independent ubiquitination activities (Hills-Muckey et al., 2022; Negishi et al., 2022).

A second TIR1 variant has been developed using a distinct mutation at the same TIR1[F79] residue: F79A (*Os*TIR1[F74A]) to increase TIR1 affinity to the synthetic auxin analog 5-adamantyl-IAA (5-Ad-IAA). This variant was named “super-sensitive AID (ssAID)” and can be activated by 5-Ad-IAA at 1000-to 10000-fold lower concentrations compared to those required of IAA for TIR1[F79], depending on the cell type (Nishimura et al., 2020; Yamada et al., 2018). Unlike F79G, F79A has not yet been adapted for use in *C. elegans*.

The AID system is now widely used across eukaryotic systems but has seen particularly rapid adoption in *C. elegans* research. The relative speed and efficiency of CRISPR and MosSCI transgenesis in *C. elegans* support a diverse set of innovations including the adoption of synthetic auxin analogues such as 1-naphthaleneacetic acid (NAA) that show enhanced solubility and photostability compared to IAA (Martinez et al., 2020), the development of a variety of single-copy, tissue-specific TIR1-expessing strains (Ashley et al., 2021); and the use of single or multiple mIAA7 epitopes as an alternative degron sequence for enhanced depletion kinetics (Sepers et al., 2022).

Though most technical development of the AID system has focused on obtaining rapid and complete degradation to replicate classical loss-of-function mutations, the AID system allows much more finesse. For example, quantitative depletion of the RNA Polymerase II (RNAPolII) subunit RPB-2 revealed a dose-dependent relationship between RNAPolII abundance and lifespan, providing in effect a quantitative dial on the rate of aging (Oswal et al., 2022). Here, we further explore the AID system as a way to provide quantitative perturbations to organismal physiology, and in doing so, reveal the major factors contributing to AID-mediated protein degradation kinetics. We adapt for the first time the TIR1[F79A] (ssAID) variant in *C. elegans*, and demonstrate that it has slightly better quantitative performance than the TIR1[F79G] (AID2) variant. Then, taking advantage of the orthogonality of the AID and AID2 systems, we demonstrate a dual-channel AID system that can independently deplete the target of interest in two different tissues by varying the concentrations of the respective TIR1 variant ligands. Finally, we re-engineered the TIR1 transgene to facilitate robust protein depletion in both the soma and germline. Taken all together, we describe the differences in protein depletion kinetics between the three versions of the AID system and establish that its components can be modulated to achieve the desired kinetics.

## RESULTS

### The different versions of the AID system have similar, but not identical performance

First, we developed a CRISPR-genome editing technique that rapidly allows conversion between TIR1 transgenes. Three versions of the AID system currently exist based on the identity of the amino acid in position 79 located at the side wall of the auxin binding pocket of the *Arabidopsis thaliana* TIR1 (AtTIR1) protein. In wild-type TIR1, there is a phenylalanine residue at position 79 (F79). Orthogonal auxin–TIR1 pairs have been developed based on a “bump-and-hole” approach where the side wall of TIR1 is carved by substituting F79 with an aliphatic residue while reciprocally introducing a bump to the indole ring of IAA to fill the cavity. In the AID2 system, F79 is substituted with glycine (F79G) and is paired with its high-affinity ligand 5-Ph-IAA. In contrast, F79 is substituted with alanine (F79A) in the ssAID system and is paired with its high-affinity ligand 5-Ad-IAA (Fig. 1b). To allow rapid conversion among these TIR1 variants, we designed a CRISPR-Cas9 strategy to edit the F79 locus of TIR1 transgenes *in vivo* (Supp. Fig. 1) to obtain three identical *C. elegans* strains differing only in a single amino acid residue. We obtained editing efficiencies of 51.4% for F79 to F79A conversion and 45.9% for F79 to F79G conversion from 37 genotyped animals each, with some homozygous conversions occurring in the first filial generation (F_1_). We conclude that our method allows rapid conversion between different engineered TIR1 alleles, allowing users to select the correct one for their purpose.

To compare the quantitative performance of these TIR1 variants in a physiologically relevant context, we chose the *daf-2* insulin/IGF receptor as a degradation target. *daf-2* is a highly pleiotropic gene with a quantitative influence on multiple phenotypes. Mutations in the insulin receptor can extend lifespan up to 300% (Gems et al., 1998), a phenotype recently recapitulated using the AID system (Venz et al., 2021). Mutations in *daf-2* are also known to influence a complex organismal decision made by *C. elegans* during development, in which a variety of sensory cues are integrated to decide whether or not to enter a spore-like diapause state (Kimura et al., 1997). Therefore, choosing *daf-2* as a target with which to engineer the AID system provides us with multiple quantitative phenotypes through which we can quantify AID degradation rates.

First, we focused on the lifespan phenotype, which allows us to quantify the integrated effect of TIR1 activity over several weeks. In all three systems, we observed a dose-dependent increase in lifespan extension, identifying a range in which the kinetics of protein depletion is rate-limited by auxin ligand concentration (Fig. 1c). At the highest concentrations of ligands tested (750 µM K-NAA, 1.875 µM 5-Ph-IAA and 5-Ad-IAA), we observed that the original F79 TIR1 variant was able to produce the greatest mean lifespan (26.5 days), followed by F79A (22.8 days), and finally, F79G (19.3 days) (Fig. 1d).

To better understand the kinetics of protein depletion, we fit hill functions to our dose-response data. This allowed us to estimate the maximum lifespan extension at saturating ligand concentrations for each TIR1 variant: 219% in F79, 203% in F79A, and 184% in F79G (Fig. 1e; dotted lines). These results again suggest that the F79 variant produces the greatest degree of DAF-2 degradation among the three variants.

To compare the cross-reactivity of the three ligands across the three TIR1 variants, we turned to the dauer assay, which allows us to quantify DAF-2 degradation during development instead of adulthood. In between the first and second larval molts, DAF-2 receptor activity determines the probability of an individual entering diapause, a morphologically distinct state we can identify visually and use as a quantitative readout of TIR1 activity. We find that much lower concentrations of ligand are required to shift animals from 0% to 100% dauer entry compared to the doses required to extend lifespan (Fig. 1f). Again using hill fits, we estimated the ligand concentrations required to produce 50% dauer entry (EC_50_) and compared these concentrations across TIR1 variants. For TIR1[F79], the EC_50_ of this variant’s canonical ligand K-NAA was 359 nM. The EC_50_ was never reached for the other two compounds. For TIR1[F79A], the EC_50_s were 6.3nM for the canonical ligand 5-Ad-IAA, 1.6 nM for 5-Ph-IAA, and 163 uM for K-NAA. Finally, for TIR1[F79G] the EC_50_s were 8.4nM for the canonical ligand 5-Ph-IAA, 48nM for 5-Ad-IAA, and the EC_50_ was never reached for K-NAA. From these data, we conclude that TIR1[F79] shows negligible cross-reactivity with 5-Ad-IAA and 5-Ph-IAA at their typical concentrations, that TIR1[F79A] cross reacts with all three ligands at their typical concentrations, and that TIR1[F79G] cross reacts with 5-Ad-IAA but not K-NAA at their typical concentrations.

### TIR1 protein abundance and per-molecule TIR1 ubiquitination activity rates are the major determinants of AID system performance

We then applied an interventional approach to identify the biochemical mechanisms determining AID system performance. We experimentally modulated two aspects of the system: first, we introduced a 3x repetition of the degron sequence on the *daf-*2 receptor (Fig. 2a). Our version of the 3x degron tag uses three copies of AID*, in contrast to the 3x mIAA7 tag previously found to improve auxin-dependent degradation at the expense of increased basal degradation (Sepers et al., 2022). Second, we altered the abundance of the TIR1 protein by replacing the somatic *eft-3* promoter with an alternative that produces lower somatic expression: the *eif-3.B* promoter. In contrast to *eft-3p*::TIR1 (Fig. 2b), the *eif-3.Bp*::TIR1 line notably shows germline expression (Fig. 2c). Having made these constructs, we then explored several permutations: pairing both 1xAID and 3xAID degron tags with the F79, F79A and F79G TIR1 variants.

**Figure 2.**
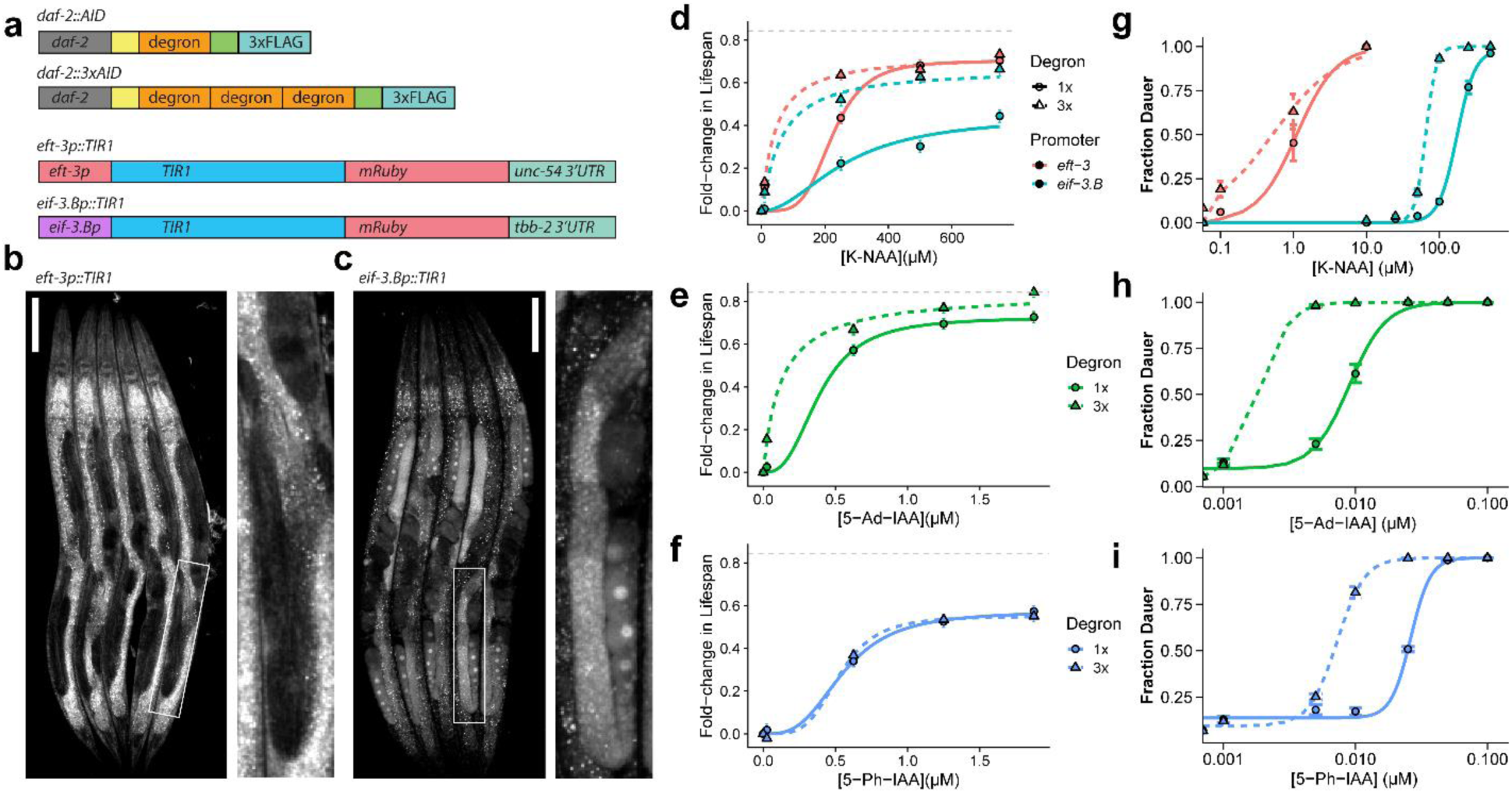
Engineering better quantitative control of the auxin-inducible degradation (AID) system. (a) Fine-tuning the AID system by modulating the number of degron repeats on the AID tag (top) or modulating TIR1 expression levels (bottom) using the *eft-3* promoter (high somatic expression, no germline expression) or the *eif-3.B* promoter (intermediate somatic expression, high germline expression). (b) Organismal expression of *eft-3p*::TIR1::mRuby, with germline (inset) expression absent. (c) Organismal expression of *eif-3.Bp*::TIR1::mRuby, with the germline (inset) expression present. (d) The dose-response effect on lifespan of K-NAA on populations expressing TIR1[F79] under the *eif-3.B* promoter (blue), *eft-3* promoter (red), combined with DAF-2::AID (solid lines; circles) or DAF-2::3xAID (dotted lines; triangles). (e) The dose response of 5-Ad-IAA on populations expressing *eft-3p*::TIR1[F79A] combined with DAF-2:AID (solid lines; circles) or DAF-2::3xAID (dotted lines; triangles). (f) The dose response of 5-Ph-IAA on populations expressing *eft-3p*::TIR1[F79G] combined with DAF-2::AID (solid lines; circles) or DAF-2::3xAID (dotted lines; triangles). (g–i) The dose response of each of the compounds and TIR1 variations in d–f, but this time characterizing the probability of dauer entry.

We then quantified the effect of these modifications to the AID system using lifespan as a readout of DAF-2::AID degradation rates. For the wild-type TIR1[F79] construct, we find that with the standard 1xAID tag, lowering the expression of TIR1 using *eif-3.B*p dramatically reduces the maximum lifespan extension obtainable (Fig. 2d). By adding a 3xAID tag, we can mostly recover the original lifespan extension. In contrast, adding a 3xAID tag to the highly-expressed *eft-3p*::TIR1 line produced only small additional lifespan extensions. These data reveal important biochemical determinants of the AID system: at saturating ligand concentrations, *eft-3p*::TIR1 is sufficiently abundant to maximally ubiquitinate DAF-2::AID and thereby maximize degradation rates with 1 degron repeat. However, after lowering TIR1 abundance using *eif-3.Bp*::TIR1, even at saturating K-NAA concentrations, the TIR1 pool available is not sufficient to maximally ubiquitinate DAF-2::AID. Then, our data demonstrates that adding 3xAID repeats to DAF-2::AID increases degradation rates, allowing each molecule of TIR1 to effect a greater degradation rate presumably through cooperative interactions among the additional ubiquitination sites available on the 3xAID.

In contrast, pairing the 3xAID tag with the TIR1[F79A] mutation substantially increased the lifespan extension produced even when the TIR1 protein was expressed at high abundance under the *eft-3* promoter. This suggests that the rate-limiting factors for TIR1[F79A]-mediated DAF-2::AID degradation are different from those for wild-type TIR1[F79]. The lower lifespan extension of TIR1[F79A] compared to TIR1[F79] in the 1xAID background highlights a lower intrinsic ubiquitination rate for each molecule of TIR1[F79A] compared to TIR1[F79], since the former fails to maximize ubiquitination rates despite sufficient TIR1 abundance. However, the intrinsic lower activity of F79A is compensated for by pairing it with a 3x degron, and we then observe the largest lifespan extension produced by any combination of TIR1 variants and degron tags. In contrast, pairing the 3xAID tag with TIR1[F79G] produced no effect on lifespan compared to 1xAID. These data suggest that the intrinsic activity of TIR1[F79G] is not only lower than TIR1[F79] and TIR1[F79A], but also that this defect cannot be compensated for by adding additional ubiquitination sites.

Finally, we compared all combinations of TIR1 and degron tags using the dauer assay. For all strains, we were again able to obtain 100% dauer entry rates (Fig. 2g; Supp. Fig 2). In concordance with our lifespan data, lowering the expression of TIR1[F79] using the *eif-3.B* promoter decreased the apparent degradation rate of DAF-2::AID, increasing the EC_50_ 150-fold from 1.1 µM to 157 µM and again highlighting the crucial dependence of AID system activity on TIR1 expression levels. Adding a 3x degron recovered some of this activity, lowering the EC_50_ by about 40%. This lowering of the EC_50_ was observed both in *eft-3p*::TIR1 and *eif-3.Bp*::TIR1, suggesting that during development, *eft-3p*::TIR1 is not expressed at sufficiently high concentrations to produce maximal ubiquitination rates with 1xAID.

Adding 3x degrons to the TIR1[F79A] and TIR1[F79G] variants again produced very large decreases in the EC_50_ of their respective ligands—much larger than the effect of 3x degron for the TIR1[F79] strain (Fig. 2h,i). These data support our model that TIR1[F79A] and TIR1[F79G] produce lower per-molecule ubiquitination rates of DAF-2, deficits that can be compensated for by increasing the availability of ubiquitination sites. In the lifespan assay, we had found that adding a 3xAID tag produced little to no effect on TIR1[F79G] activity, but in the dauer assay, the effects we see from adding the 3xAID tag suggest that adding additional ubiquitination sites can improve TIR1[F79G] activity when TIR1[F79G] is expressed at lower than saturating abundances—a situation we infer is the case during development but not during adulthood.

In conclusion, our data reveals important determinants of AID activity, most notably the requirement for TIR1 to be expressed at high abundance relative to degradation targets to ensure TIR1 abundance is not the limiting factor for degradation rates. Regardless of other benefits of the TIR1[F79A] and TIR1[F79G] variants, we find that these two proteins exhibit substantially lower overall TIR1 ubiquitination activity per molecule, to the extent that TIR1 abundance is limiting even at very high expression levels. For TIR1[F79A] but not TIR1[F79G], this deficit can be ameliorated by adding a 3xAID tag to the degradation target.

### Transcriptomic analysis confirms the relative activities across TIR1 variants

We then sought to develop a third quantitative assay to study the quantitative and qualitative differences in TIR1 function across the different TIR1 variants, promoters, and degron tags. Because *daf-2* mutations produce strong effects on organismal gene expression (Murphy et al., 2003), we set out to study AID performance by collecting whole animal transcriptomes from the eight TIR1-expressing strains considered so far: *eft-3p*::TIR1[F79], *eif-3.Bp*::TIR1[F79], *eft-3p*::TIR1[F79A], and *eft-3p*::TIR1[F79G] each combined with either DAF-2::AID or DAF-2::3xAID— with and without 24 hours of exposure to the highest concentration of their respective ligands, K-NAA (750 µM) and 5-Ph-IAA or 5-Ad-IAA (1.875 µM).

Principal component analysis (PCA) revealed strong effects of the AID system on gene expression (Fig. 3a). The first principal component (PC), explaining 52% of all inter-sample variation (Supp. Fig. 3a), reveals a clear separation between strains expected to degrade DAF-2::AID (high PC1 values) from the remainder: the strains unexposed to auxin along with auxin-exposed strains lacking the TIR1 transgene exhibited low PC1 values, in contrast to all eight strains with TIR1 transgenes exposed to auxin which exhibited high PC1 values. Therefore, we conclude that most of the gene regulatory differences between strains results from the influence of TIR1-mediated DAF-2 degradation on gene expression.

**Figure 3.**
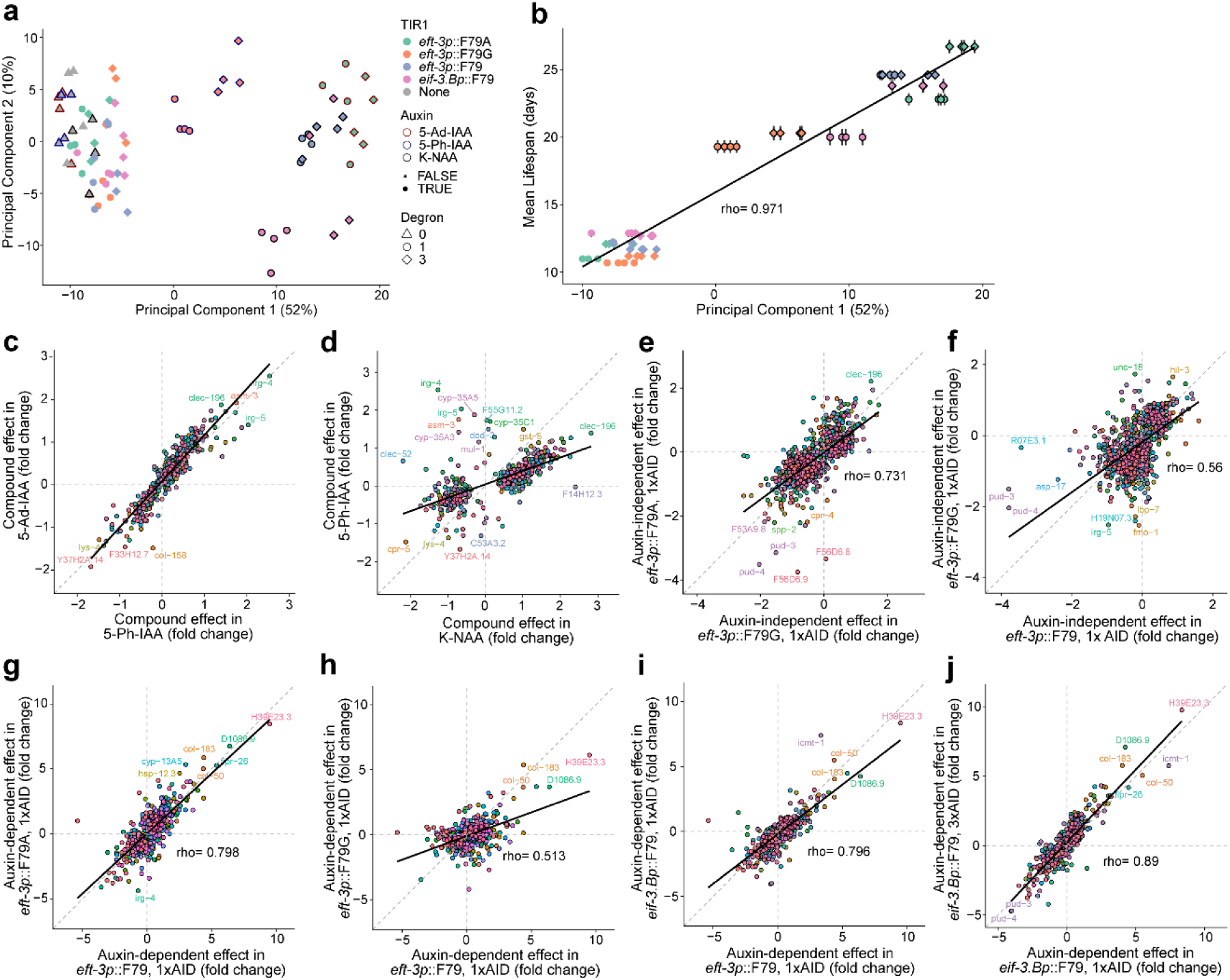
The independent effects of AID system components on gene-expression and physiology. mRNA was collected on the first day of adulthood from wild-type populations (gray), eight TIR1-expressing strains: *eft-3p*::TIR1[F79] (blue) and *eif-3.Bp*::TIR1[F79] (pink) each combined with either *daf-2*::AID (circles) or *daf-2*::3xAID (diamonds) tags, and also *eft-3p*::TIR1[F79A] (green) and *eft-3p*::TIR1[F79G] (orange) strains, as well as a wild-type population with no genome edits (gray). Populations were exposed to no compound (no edge), K-NAA (black edge), 5-Ph-IAA (blue edge), or 5-Ad-IAA (red edge). (a) Principal Component (PC) analysis of all strains was performed, and the projection of each population’s transcriptome was plotted against PC1 and PC2. (b) The same PC1 projection of each population compared to the mean lifespan of each strain, with linear fit (line). (c) The effect on gene expression of exposing wild-type animals to 5-Ph-IAA, compared to the effect of 5-Ad-IAA, with linear regression (black). (d) The effect of exposing wild-type animals to K-NAA, compared to the effect of 5-Ph-IAA. (e) In the absence of any compound in *daf-2*::AID populations, the effect of *eft-3p*::TIR1[F79G] transgene on gene expression compared to the effect of *eft-3p*::TIR1[F79A]. (f) In the absence of any compound, the effect of *eft-3p*::TIR1[F79] compared to *eft-3p*::TIR1[F79G]. Fold-changes calculated for each strain relative to wild-type populations expressing no transgenes. (g) In the presence of their respective activating compounds, the effect on gene expression of TIR1[F79] compared to TIR1[F79A]. (h) In the presence of their respective activating compounds, the effect on gene-expression of TIR1[F79] compared to TIR1[F79G]. (i) In the presence of 750 µM K-NAA in *daf-2*::AID populations, the effect on gene expression of TIR1 expressed under the *eft-3* (x-axis) vs. the *eif-3.B* promoter (y-axis). (j) In the presence of 750 µM K-NAA in *eif-3.Bp*::TIR1[F79] populations, the effect on gene expression of adding the *daf-2*::1xAID (x-axis) vs. *daf-2*::3xAID (y-axis). Fold-changes calculated for each strain relative to wild-type populations expressing no transgenes exposed to the same compound.

Remarkably, we find that PC1 captures an aspect of gene expression on day 2 that is highly correlated with each strain’s remaining lifespan (rho = 0.971; Fig. 3b; Supp. Fig 3a) demonstrating a strong relationship between the strength of gene expression changes produced by auxin-induced DAF-2 degradation measured on day 2 with the physiologic consequences of such degradation on aging, measured weeks later. This high degree of correlation is maintained when considering only auxin-dependent effects (Supp. Fig. 3c) after segregating the auxin-independent (Supp. Fig. 3d).

Though we did not identify any effects of auxin ligands on lifespan or dauer entry in the absence of TIR1 transgenes, we nevertheless considered the possibility that exposure to auxin ligands alone might have an effect on gene expression. In wild-type populations, we find that hundreds of gene regulatory changes are produced by exposure to 750uM K-NAA, 1.875uM 5-Ph-IAA, and 1.875uM 5-Ad-IAA (Supp. Table 1). The effects are strongly correlated across all three ligands, with 5-Ph-IAA and 5-Ad-IAA producing practically identical effects (Fig. 3c). However, the magnitude of gene-expression changes produced by these two compounds was substantially lower than by K-NAA (linear fit, slope << 1; Fig. 3d, Supp. Fig. 3e), perhaps reflecting the much higher concentration of K-NAA used. However, there were a small number of genes that showed qualitatively distinct differential regulation by K-NAA compared to 5-Ph-IAA and 5-Ad-IAA, suggesting some distinct physiological effects of K-NAA compared to the other two.

### The physiological influence of auxin-like ligands

There are many reports in the literature that TIR1 transgenes can influence physiology even in the absence of auxin, a phenomenon called “basal activity”. We investigated basal activity by considering the gene regulatory effect of each TIR1 variant in the presence of DAF-2::AID but in the absence of any auxin ligand. We find a large number of differentially regulated transcripts in each TIR1 strain compared to wildtype, with the basal activity of F79G and F79A variants producing highly correlated (rho=.731) effects (Fig. 3e) and F79 producing less strongly correlated effects (rho=.56) compared to the other two (Fig. 3f, Supp. Fig. 3f). We conclude that our gene expression experiments highlight an auxin ligand-independent activity of all TIR1 variants.

### Quantitative and qualitative differences between TIR1 variants

Finally, we investigated the major gene regulatory effects identified by our study—the auxin ligand-dependent, TIR1-dependent effects of DAF-2 degradation. We found a large number of strongly differentially regulated genes (Supp. Table 2), this time strongly correlated between F79 and F79A (rho=0.798, Fig. 3g) and to a lesser extent between F79G and the other two (Fig. 3h, rho=0.51; Supp. Fig. 3g, rho=0.667). The magnitude of gene expression changes produced by F79G was less than that of F79 (linear fit slope 1) and F79A (linear fit slope 1), consistent with the greater effect of the F79 and F79A variants on lifespan compared to F79G (Fig. 1e).

We then considered the effect of TIR1 abundance on gene expression, comparing the effect of auxin-induced DAF-2 knockdown in *eft-3p*::TIR1 and *eif-3.B*p::TIR1. Remarkably, despite the differences in both expression level and localization, we see a strong correlation in the effects on gene expression in both strains (rho=0.796), though these effects are of smaller magnitude in the less abundant *eif-3.Bp*::TIR1 (slope < 1, Figure 3i). We see an even higher correlation (rho=.89) between the effects of *eif-3.Bp*::TIR1 in the presence of 1xAID compared to 3xAID (Fig. 3j) as well as in the *eft-3p*::TIR1[F79], eft3*-p*::TIR1[F79G] and *eft-3p*::TIR1[F79A] backgrounds (Supp. Fig. 3h–j), which was surprising given the large differences in lifespan between 1xAID and 3xAID observed at the auxin concentrations used (Fig. 2d). We conclude that the 3xAID influences lifespan through either 1) a specific set of genes that are more up-or downregulated in the 3xAID strain compared to the 1x strain; or 2) the cumulative effect of small quantitative changes across many genes that do not individually have statistical significance. As a final option, 3) the 3xAID may influence lifespan by modulating some post-transcriptional activity of DAF-2.

In conclusion, our transcriptomic analysis highlights an overall shared influence on gene expression of the different TIR1 variants driven by various promoters when paired with the 1x and 3xAID tags. Such similarity is to be expected by the shared molecular target in every case: the DAF-2::AID degron. However, our findings also show some off-target effects produced by the different auxin ligands, as well as some differences in the gene expression effects of DAF-2::AID degradation when produced by different variants: TIR1[F79] and TIR1[F79A] show nearly identical auxin-dependent effects on gene expression that differ slightly from TIR1[F79G].

### A “dual-channel” AID system combines orthogonal, tissue-specific TIR1 variants

A major constraint of the auxin-AID system is that it allows degradation of only one target at a time. However, our data (Fig. 1f) highlights an orthogonality between different TIR1 variants that we might use to create a “dual-channel” AID system to enable independent depletion of proteins in two different tissues. Since concentration regimes exist in which TIR1[F79] and TIR1[F79G] can be independently activated by K-NAA and 5-Ph-IAA, respectively, with no cross-talk (Nishimura et al., 2020; Uchida et al., 2018; Yesbolatova et al., 2020) (Fig. 1f), we reasoned that both variants could be expressed in non-overlapping tissues to allow independent quantitative control over a protein’s degradation rate in different tissues. As a proof-of-concept, we decided to express the TIR1[F79] and TIR1[F79G] variants using intestinal and neuronal-specific promoters, respectively (Fig. 4a). The neuronal-intestinal axis is important in many biological processes in *C. elegans* including aging (Morphis et al., 2022; Roy et al., 2022; Uno et al., 2021; B. Zhang et al., 2018; Y. P. Zhang et al., 2022), chemosensory behavior (Lee & Mylonakis, 2017; Matty et al., 2022), rhythmic behavior (Mahoney et al., 2008), and innate immunity (Foster et al., 2020; Liu et al., 2023; Nag et al., 2017).

**Figure 4.**
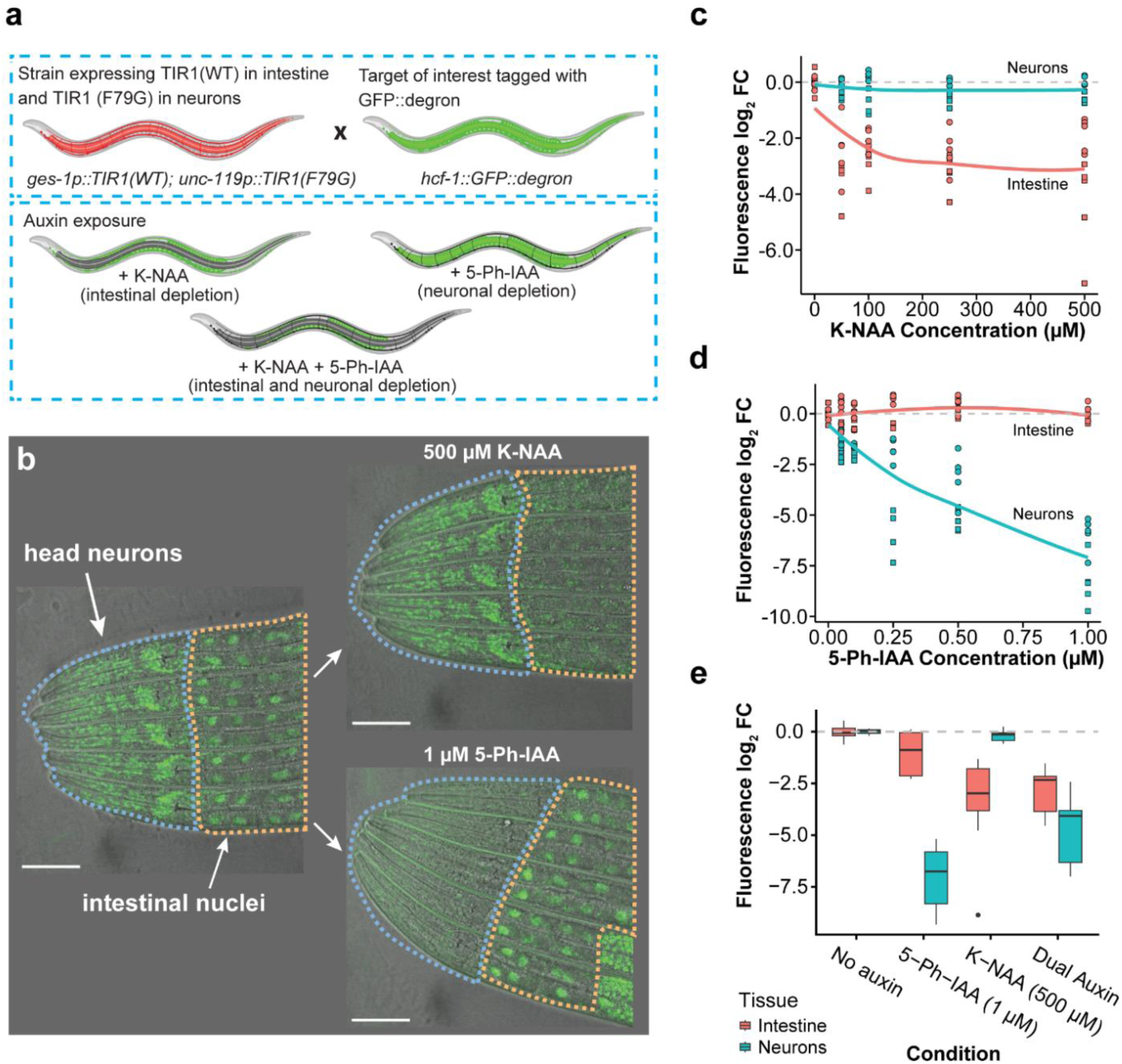
A tissue-specific “dual-channel” AID system. The dual-channel AID system results in tissue-specific degradation of the target of interest. (a) A strain harboring the HCF-1::GFP::degron reporter was crossed with a strain containing two tissue-specific TIR1 transgenes: *ges-1p*::TIR1 (intestine) and *unc-119p*::TIR1[F79G]. Depending on the auxin analogue added, HCF-1::GFP can be selectively depleted in the intestine or neurons, or can be simultaneously depleted in both tissues. (b) Representative image of dual-channel TIR1, HCF-1::GFP::degron worms without auxin shows localization of GFP signal in neuronal and intestinal nuclei (left panel). Addition of 500 µM K-NAA results in degradation of the target in the intestine (upper right panel) while addition of 1 µM 5-Ph-IAA results in degradation of the target in neurons (lower right panel). (c) Exposure to varying concentrations of K-NAA shows preference for degradation of the target in the intestine. (d) Exposure to 5-Ph-IAA results in the opposite effect, with preferential degradation of the target occurring in neurons. (e) Exposure to both K-NAA and 5-Ph-IAA results in simultaneous depletion in both intestine and neurons. Fluorescence log_2_ FC: log_2_ fold change of fluorescent signal quantified using absolute photon counts normalized to the no auxin condition. Points represent the signal quantified from individual worms and the shape of the points corresponds to each independent replicate (N = 2).

We then chose HCF-1::GFP::AID as a brightly-expressed degradation target and quantitative readout of TIR1 activity. With the combination of TIR1 variants used, we hypothesized that organismal exposure to K-NAA should deplete HCF-1 specifically in the intestine while organismal exposure to 5-Ph-IAA should deplete HCF-1 specifically in neurons (Fig. 4a). Using fluorescence as a measure of HCF-1 abundance, we observed exactly this effect (Fig. 4b). Across replicates, we observed a dose-dependent degradation of intestinal HCF-1 with respect to K-NAA (Fig. 4c) and a dose-dependent degradation of neuronal HCF-1 in response to 5-Ph-IAA (Fig. 4d). When individuals were exposed to both compounds simultaneously, we observed degradation in both neurons and the intestine (Fig. 4e), though neuronal degradation rates appeared to be lowered slightly in the presence of both compounds compared to only 5-Ph-IAA, perhaps through the competitive occupation of TIR1[F79G] binding sites by K-NAA. Based on these results, we conclude that a combination of TIR1[F79] and TIR1[F79G] with distinct tissue-specific expression patterns can be used to obtain independent control of target protein levels simultaneously in two different tissues.

### Transgene de-silencing complicates design of a germline-soma dual-channel AID system

We then sought to generate a “dual-channel” system that would allow independent degradation of proteins in the soma and the germline. To do this, we combined the pan-somatic *eft-3p*::TIR1::*unc-54* 3’UTR (L. Zhang et al., 2015) with a newly created germline-expressed *mex-5p*::TIR1[F79G]::*tbb-2* 3’UTR transgene. To visualize AID activity across all tissues, we chose RPB-2 (Oswal et al., 2022), the second-largest subunit of RNA polymerase II, as a bright nuclear-localized protein visible both in somatic and germline nuclei. Surprisingly, this combination of TIR1 constructs did not produce the expected tissue-specific responses (**Supp. Fig. 4a**), as exposure to K-NAA resulted in degradation of RPB-2::AID both in the soma (**Supp. Fig. 4b**) and in the germline (**Supp. Fig 4c**). Remarkably, we found that the germline depletion of RPB-2::AID required both somatic *eft-3p::TIR1* and germ line *mex-5p::*TIR1[F79G] transgenes, as no germline depletion was observed when either TIR1 transgene was present in isolation (**Supp. Fig. 4c**). We therefore arrive at the surprising hypothesis that expression of the K-NAA-insensitive *mex-5p*::TIR1[F79G] transgene must somehow enable the K-NAA-sensitive pan-somatic *eft-3p*::TIR1[F79] transgene. Using confocal microscopy, we find that *eft-3p*::TIR1::mRuby gains germline expression when co-expressed with the *mex-5p*::TIR1[F79G] transgene (**Supp. Fig. 4d**). We therefore explain the surprising failure of the germline-soma dual channel AID construct: activation of germline *eft-3p*::TIR1 expression by *mex-5p*::TIR1[F79G] allows KNAA to deplete RBP-2::AID both in the soma and the germline.

A major determinant of germline transgene expression is gene silencing mediated by PIWI-interacting RNAs (piRNAs) (Ashe et al., 2012; Shirayama et al., 2012). We therefore investigated whether presence of the germline-licensed *mex-5p*::TIR1[F79G] might result in the de-silencing of the nearly-homologous *eft-3p*::TIR1 transgene in the germline (Shirayama et al., 2012). We disrupted HRDE-1 via RNAi to globally inhibit germline silencing (Ding et al., 2023), and after five generations of *hrde-1* RNAi we began to detect *eft-3p*::TIR1 expression in the germline, even in the absence of *mex-5p*::TIR1 (**Supp. Fig. 5a**).We confirmed that de-silencing was sufficient to enable K-NAA-dependent RPB-2::GFP degradation (**Supp. Fig. 5b-d**). From these data, we conclude that piRNA-mediated transgene silencing is required for the pan-somatic expression pattern of *eft-3p::*TIR1 and that disruption of this silencing is sufficient to convert *eft-3p::*TIR1 into whole-body expression pattern. We therefore conclude that a germline/soma dual-channel auxin degron system is not possible using existing TIR1 transgenes, as the required pan-somatic promoter—a promoter whose exclusion from the germline does not depend on germline silencing—has not yet been developed.

### Pan-organismal knockdown of proteins using a re-engineered TIR1

Our better understanding of *eft-3*p::TIR1 suggested a path for producing a true pan-organismal auxin-inducible degron system. Such a system would allow AID experiments that recapitulate the effect of conventional mutations and RNAi experiments which typically involve knockdown of protein targets across all tissues. We hypothesized that such a pan-organismal AID system could be obtained by creating a new *eft-3p*::TIR1 construct engineered to circumvent piRNA activity (Aljohani et al., 2020). To obtain such a construct, we sought to eliminate the germline silencing of the TIR1 transgene by including an early first intron, the endogenous Periodic A_n_/T_n_ Clusters (PATCs), and using the *gpd-2/3* intergenic sequence for co-expression (Aljohani et al., 2020) (**Figure 5a**). In addition, we codon-optimized the entire TIR1 coding sequence and removed potential piRNA-binding sites. We chose to swap the *unc-54* 3’UTR to the β-tubulin 3’UTR (*tbb-2*) as the latter has been described as enabling expression in all germ cells (Merritt et al., 2008).

**Figure 5.**
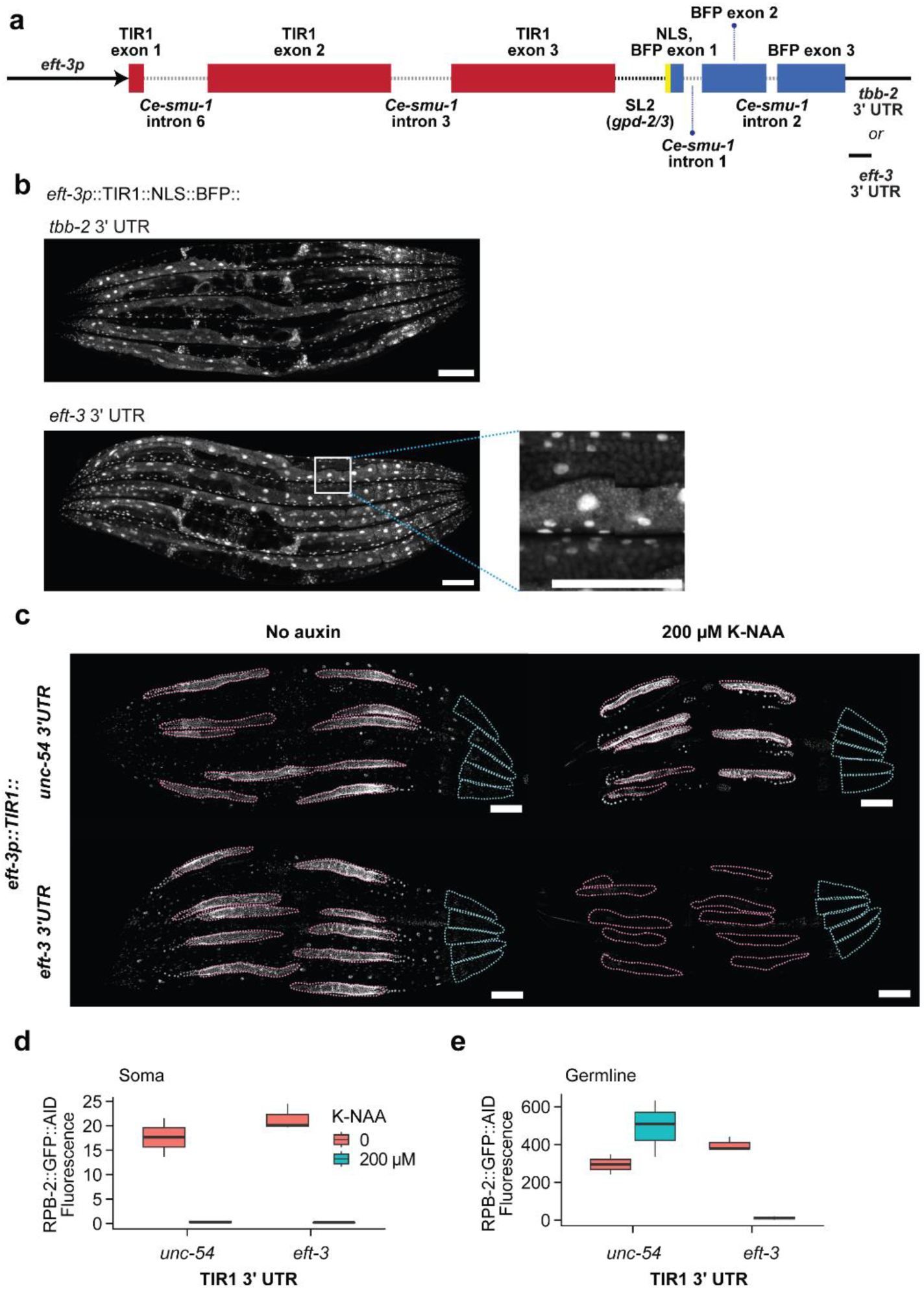
An improved whole-organism AID degradation system. A re-engineered TIR1 transgene with somatic and germline expression is capable of depleting target proteins in the whole body. (a) Schematic of the redesigned TIR1 transgene consisting of the *eft-3* promoter, TIR1 with *smu-1* introns, SL2 trans-splicing sequence from the *gpd-2/3* operon, c-Myc nuclear localization signal (NLS), mTagBFP2 with *smu-1* introns, and a *tbb-2* or *eft-3* 3’UTR. The diagram is drawn to scale. (b) Confocal images of the redesigned TIR1 transgenes containing either the *tbb-2* 3’UTR or the *eft-3* 3’UTR. The inset highlights the expression of TIR1::mTagBFP2 in the germline of *eft-3* 3’UTR animals. (c) Comparison of somatic and germline protein depletion of degron-tagged RPB-2::EGFP between *eft-3p*::TIR1::mRuby::*unc-54* 3’UTR and our redesigned *eft-3p*::TIR1::mTagBFP2::*eft-3* 3’UTR transgene upon auxin exposure. (d) Quantification of somatic and (e) germline RPB-2::GFP signal in *eft-3p*::TIR1::*unc-54* 3’UTR and *eft-3p*::TIR1::*eft-3* 3’UTR animals on and off auxin. Data for (d) and (e) are derived from five worms per condition from a single biological replicate.

Using MosTI, we successfully integrated a single copy of *eft-3p*::TIR1::SL2::NLS::BFP:: *tbb-2* 3’UTR. However, this transgene was not expressed in the germline (**Figure 5b**). Exposure to *hrde-1* RNAi for several generations again resulted in de-silencing of germline TIR1 expression (**Supp. Fig. 5e**), demonstrating that our changes did not eliminate piRNA-mediated transgene silencing. Since the endogenous EFT-3 protein is expressed in the germline, we hypothesized that germline expression of TIR1 might be rescued by switching from the *tbb-2* 3’UTR to the native *eft-3* 3’UTR. We therefore performed a 3’UTR swap using CRISPR–Cas9, excising the existing *tbb-2* 3’UTR from the TIR1 transgene and replacing it with the native *eft-3* 3’UTR sequence. Happily, we find this final modification was sufficient to enable germline expression of TIR1, providing the pan-organismal TIR1 transgene: *eft-3p*::TIR1::SL2::NLS::BFP::*eft-3* 3’UTR (**Figure 5b**). To confirm that this transgene exhibits TIR1 activity in both somatic tissues and the germline, we returned to the degron-tagged RPB-2::GFP as a reporter for AID activity. In contrast to the *eft-3p*::TIR1::*unc-54* 3’UTR precursor, the *eft-3p::*TIR1::*eft-3* 3’UTR transgene degrades RPB-2::AID across all tissues (**Figure 5c**). We conclude that our investigation into piRNA-mediated transgene silencing in the AID system allowed us to generate a new pan-organismal TIR1 transgene.

## DISCUSSION

In this work, we applied engineering approaches to identify the biochemical factors determining the performance of the AID system in the multicellular animal *C. elegans*. We find that TIR1 abundance and ubiquitin binding site availability interact to determine the chemical kinetics of the auxin-inducible degron system. Notably, our analysis identifies a lower per-molecule activity of both TIR1[F79A] (ssAID) and TIR1[F79G] (AID2) compared to the wild-type TIR1[F79] (AID) variant. In some cases, these deficiencies can be ameliorated by increasing ubiquitination sites using 3x degron tags. With 3x degron tags, we identified the best-performing AID system as the TIR1[F79A] variant, introduced into *C. elegans* for the first time here.

We then applied our quantitative understanding of the different AID variants to design and implement a “dual-channel” AID system that combines TIR1[F79] and TIR1[F79G] variants expressed in distinct tissues to effect simultaneous, independent control of target protein abundance in a tissue-specific manner. In multicellular animals, single proteins often play distinct physiological roles in different tissues (Jeffery, 1999; Wang et al., 2008; Yang et al., 2016), and our technique provides a new approach for interventional investigations into such phenomena by quantitatively manipulating the abundance of a single protein in two distinct tissues. However, our “dual-channel” technique can be applied, with no additional modification, to independently manipulate two separate degradation targets—in the special case where the two targets exhibit non-overlapping tissue expressions. For example, had we tagged two proteins with AID tags, first a neuron-specific, and then second, an intestinal-specific protein, our dual-channel approach using TIR[F79] and TIR[F79G] would provide simultaneous, independent manipulation of the two separate tissue-specific proteins.

Finally, we generated a re-engineered TIR1 transgene capable of robustly depleting target proteins in both the soma and the germline. In total, we demonstrate how a better quantitative understanding of the auxin-inducible degron system can be used to design technologies that support advanced experimental interventions into organismal physiology.

## MATERIALS AND METHODS

### *C. elegans* strains and maintenance

*Caenorhabditis elegans* strains were maintained on Nematode Growth Medium (NGM) plates seeded with *Escherichia coli* OP50 bacteria (Stiernagle, 2006). Strains were maintained at 20°C, except in lifespan assays where worms were kept at 25°C. The names and genotypes of all strains generated in this study are listed in Supplementary Table 3.

### Auxin experiments

#### Plate preparation

Potassium 1-Naphthaleneacetate (K-NAA) and 5-Adamantyl-IAA (5-Ad-IAA) were obtained from TCI (N0006 and A3390, respectively) while 5-Ph-IAA was obtained from MedChemExpress (HY-134653). For K-NAA, a 500 mM stock solution was made by dissolving the salt in M9 buffer and stored at −20°C. For 5-Ad-IAA and 5-Ph-IAA, 5 mM stock solutions were made by dissolving the salt in DMSO and stored at −80°C. Before addition to molten NGM, the stock solutions were further diluted in M9 depending on the final concentration required. Molten NGM was kept at a temperature of 56°C using a water bath prior to auxin supplementation and plate pouring. Plates were seeded with fresh *E. coli* OP50.

#### Dauer and lifespan assays

For Dauer and lifespan assays, DAF-2::1x degron or DAF-2::3x degron embryos were synchronized by bleaching gravid hermaphrodites. For Dauer assays, approximately 150 eggs were placed per 55-mm plate immediately after bleaching and were maintained at 20°C. Scoring for Dauer formation was performed after 48 hours. Three plates were scored per condition, in three independent trials (N∼2000 worms per condition). For lifespan assays, synchronized embryos were allowed to grow on NGM plates without auxin for 48 hours at 25°C (up to the late L4 stage), before being transferred to auxin plates supplemented with 5-Fluoro-2’-deoxyuridine (FUDR, Sigma-Aldrich, F0503) at a final concentration of 40 µM. Approximately 40 worms were placed per 55-mm plate, with four plates per condition. Experiments in the TIR1[WT] background was performed in two independent trials (N∼320 worms per condition) while experiments in the TIR1[F79G] and TIR1[F79A] backgrounds were performed in a single trial (N∼160 worms per condition). Dead and alive worms were scored approximately every other day by gently touching the worm’s nose with the tip of a platinum wire. Worms that burst, climbed onto walls, or missing were censored from the analysis. Worms surviving to day 20 of the assay were carefully transferred to fresh auxin plates. Worms were maintained at 25°C for all lifespan assays.

#### Fluorescence assays

HCF-1::GFP and RPB-2::GFP worms were synchronized by egg layoff of gravid hermaphrodites within a two-hour window. For HCF-1::GFP experiments, worms were transferred to auxin plates at the L3 stage while for RPB-2::GFP experiments worms were transferred to auxin plates as one-day-old adults (D1). Worms were exposed to different concentrations of auxin for 24 hours prior to confocal imaging.

### Constructs and cloning

The donor vector for the single-copy integration of *eft-3p*::TIR1::SL2::NLS::mTagBFP2::*tbb-2* 3’UTR was constructed through Gibson cloning. The *eft-3p* and *tbb-2* 3’UTR fragments were amplified from *C. elegans* N2 genomic DNA while the TIR1 and SL2::NLS::mTagBFP2 fragments were ordered as gBlocks from Integrated DNA Technologies (IDT). The gBlocks were codon-optimized using IDT’s codon optimization tool. The codon-optimized sequences were then scanned for potential piRNA binding sites using PirScan (Wu et al., 2018), and silent mutations were introduced where necessary. Finally, introns from the *C. elegans smu-1* gene were introduced into the coding sequences of TIR1 (*Ce-smu-1* introns 6 and 3) and mTagBFP2 (*Ce-smu-1* introns 1 and 2). The *eft-3p*, gBlock1 (TIR1), gBlock2 (SL2::NLS::mTagBFP2), and *tbb-2* 3’UTR fragments were joint together using overlap extension PCR to form the full fragment which we designate as nTIR1. The nTIR1 fragment was cloned into the pSEM246 vector using in-house Gibson cloning reactions. pSEM246 – MCS MosTI *unc-119* target was a gift from Christian Frøkjær-Jensen (Addgene plasmid # 159821). The resulting vector was transformed into 10-beta Competent *E. coli* (High Efficiency) cells (New England Biolabs, C3019H) following the manufacturer’s protocol. Selected transformants were sequence-verified using the Plasmidsaurus whole plasmid sequencing service. The final clone harboring the TIR1 transgene and MosTI donor sequences is designated as pNES0036. All oligos and constructs are listed in Supplementary Table 4. The full nTIR1 sequence is supplied in Supplementary Table 5.

The degron 1x and 3x sequences were based on the AID* sequence, with the addition of an N-terminal GSGGGG linker. The degron sequences were codon-optimized using IDT’s codon-optimization tool and were ordered as gBlocks. The codon-optimized AID1x and AID3x sequences are listed in Supplementary Table 5.

### Transgenesis

#### daf-2::1xAID and daf-2::3xAID

The double-stranded repair template containing the 1xAID or 3xAID sequence was amplified from the corresponding gBlock using Phusion high-fidelity polymerase (Thermo Fisher Scientific, F530L). CRISPR–Cas9 transgenesis of the *daf-2* C-terminal target region was then performed following a standard microinjection protocol (Vicencio et al., 2019).

#### TIR1[F79G] and TIR1[F79A]

To generate TIR1[F79G] and TIR1[F79A], corresponding single-amino acid substitutions were introduced into the TIR1[WT] sequence in the CA1200 strain (L. Zhang et al., 2015) using CRISPR– Cas9. The single-stranded DNA (ssDNA) repair templates were ordered as 4 nmole Ultramers from IDT. F_1_ candidates were screened using amplification-refractory mutation system (ARMS) PCR (Little, 1995). To generate the germline-specific and neuronal-specific TIR1[F79G] transgenes *mex-5p*::TIR1[F79G]::F2A:mTagBFP2::AID*::NLS::*tbb-2* 3’UTR (*ohm52*) and *unc-119p*::TIR1[F79G]::mRuby (*ohm24*), respectively, the JDW221 and HAL227 strains containing the TIR1[WT] sequence were modified using CRISPR–Cas9 as previously described. JDW221 and HAL227 are gifts from Jordan Ward and Hannes Lans, respectively.

#### eif-3.Bp::TIR1

The *eif-3.Bp*::TIR1::linker::mCherry^ΔpiRNA^::*tbb-2* 3’UTR transgene was integrated into the EG6699 strain as a single-copy transgene using MosSCI (Frøkjær-Jensen et al., 2008), resulting in the JA1880 strain. The insertion was sequence-verified through genomic DNA sequencing by performing de novo alignment of reads to contigs, followed by mapping of contigs to the reference genome. Contigs that contain exogenous sequences confirm a single-copy insertion of *eif-3.Bp*::TIR1::*tbb-2* 3’UTR in the *ttTi5605* Mos1 insertion site in Chromosome II.

#### nTIR1

The redesigned *eft-3p*::nTIR1::SL2::NLS::mTagBFP2::*tbb-2* 3’UTR transgene was integrated into the CFJ94 strain as a single-copy transgene in Chromosome IV using MosTI with the *unc-119* selection approach (Mouridi et al., 2022), resulting in the AMP228 strain. The sequence of the insertion was verified using the Plasmidsaurus amplicon sequencing service. Then, the *Cbr-unc-119* rescue fragment was floxed after injection with Cre recombinase, and the resulting *unc* strains were outcrossed twice to N2 to remove the *unc-119(ed3)* allele, resulting in the AMP245 strain. The replacement of the *tbb-2* 3’UTR to the *eft-3* 3’UTR was accomplished through CRISPR– Cas9 using two guide RNAs that excised the *tbb-2* 3’UTR. The *eft-3* 3’UTR repair template was ordered as an ssDNA Ultramer from IDT. The resulting strain containing the *eft-3p*::nTIR1::SL2::NLS::mTagBFP2::*eft-3* 3’UTR *(ohm53)* transgene is designated as AMP267.

### RNAi experiments

CA1200 and AMP245 worms were cultured in NGM plates supplemented with 0.5 mg/mL ampicillin and 2 mM Isopropyl ß-D-1-thiogalactopyranoside (IPTG) seeded with HT115 empty vector (EV) or *hrde-1* RNAi from the Ahringer library (Kamath & Ahringer, 2003). Worms were maintained at 25°C and worms were transferred to fresh RNAi plates after every other generation.

### Nematode population transcriptomics

mRNA sequencing was conducted using the Smart-Seq2 technology (Picelli et al., 2014), which has been recently adapted for nematode organismal transcriptomics studies (Serra et al., 2018) and proven to be an accurate and cost-effective method for capturing population-scale gene expression (Eder et al., 2024). Lysis buffer was prepared following the published Smart-Seq2 protocols, with ERCC spike-ins (Jiang et al., 2011) added to a final dilution of 1:40,000. Thirty synchronized adult worms were individually picked into 120 µL of lysis buffer, with four replicates performed. The nematode suspensions were shock-frozen in liquid nitrogen and stored at −80°C. Lysis was carried out at 65°C for 10 minutes, followed by enzyme inactivation at 85°C for 5 minutes. cDNA libraries were then prepared according to the Smart-Seq2 protocol and purified using in-house prepared SPRI paramagnetic beads, which mimic AMPure XP beads (Beckman Coulter), with a bead-to-sample ratio of 0.8. The size distribution of the libraries was assessed using a Tapestation 4150 (Agilent), and cDNA concentration was measured using Quant-it (Invitrogen) on a plate reader (Tecan). Nextera sequencing libraries were prepared from the cDNA through tagmentation and subsequent PCR amplification with indexing primers, following the Nextera DNA library prep protocol (Illumina). These libraries were purified twice using in-house prepared SPRI paramagnetic beads with a bead-to-sample ratio of 0.9. The size distribution was again evaluated using a Tapestation 4150 (Agilent), and library concentration was determined using Quant-it (Invitrogen) on a plate reader (Tecan). The nematode RNAseq libraries were pooled in equal masses, and the paired-end Nextera libraries were sequenced on an Illumina NextSeq 500 using high-output 75-cycle v2.5 kits (Illumina), generating read lengths of 38 bases and achieving more than 2.0 x 10^6 reads per sample.

RNA-seq reads were aligned to the *C. elegans* Wormbase reference genome (release WS265) using STAR version 2.6.0c (Dobin et al., 2013), with modifications to include ERCC spike-in sequences. Gene counts were quantified with featureCounts version 2.0.0 (Liao et al., 2014). The count matrix underwent a detection threshold, requiring at least 5 counts in at least half of the samples. Genome coverage of the reads was calculated using BEDTools (Quinlan & Hall, 2010).

### Image acquisition

#### Confocal microscopy

Worms were anesthetized on 5% agarose pads using 5 mM of levamisole dissolved in M9 (ITW Reagents, P110801) and secured with a #1.5 coverslip. RPB-2::EGFP and HCF-1::EGFP images were acquired in fluorescence lifetime imaging (FLIM) mode using a Leica SP8 FALCON microscope equipped with a white light laser (WLL), a 40x Plan Apochromat, 1.3 NA oil immersion objective controlled by Leica Application Suite X software (version 3.5.7.23225). The laser was set to the 488 nm excitation line with a pulse frequency of 40 MHz and the HyD detector set to photon counting mode between the 500 to 570 nm emission spectrum. CA1200 TIR1::mRuby images were imaged using 561 nm excitation and 575 to 645 nm emission spectrum. TIR1::BFP images for RNAi experiments were acquired using a Leica SP5 inverted microscope equipped with a 405 nm diode laser, a 40x Plan Apochromat, 1.25–0.75 NA oil immersion objective controlled by LAS AF software (version 2.6.3. 8173). AMP100 and AMP281 TIR1::BFP images and TIR1::mRuby images were acquired using a Zeiss LSM 980 microscope equipped with 405 nm and 561 nm diode lasers, and a 20x Plan Apochromat, 0.8 NA dry objective controlled by Zen blue software (version 3.3.89.008).

### Image processing and analysis

GFP images acquired in FLIM mode were thresholded for photon counts in the Leica Application Suite X software. Mono-exponential reconvolution was performed by setting the lifetime to 2.6 ns. Images were then thresholded for lifetime in FIJI (version 2.14.0/1.54f), with pixels with an average lifetime less than 1.75 ns considered as autofluorescence and excluded from the analysis. Regions of interest were drawn based on brightfield images and fluorescence signals were quantified using a custom Python (version 3.12.3) script using the napari library. For HCF-1::GFP signal in the neurons, the region of interest spanned the lower half of the first bulb of the pharynx down to the end of the second bulb of the pharynx. For signal in the intestines, the region of interest covered the area demarcated by the first two pairs of anterior intestinal cells. For RPB-2::GFP images, signal in the entire head region (from the tip of the head to the end of the second pharyngeal bulb) was used as a proxy for signal in somatic cells. For signal in the germline, the region of interest covered the distal germline spanning the mitotic zone and the pachytene region before the loop. All images acquired were processed for display by cropping, rotating, and adjusting brightness and contrast using Fiji. All scale bars are set to 100 µM. No additional image manipulations were performed.

### Statistical analysis

All data plotting and statistical analysis was performed in R (version 4.2.3). Data plots were generated using the ggplot2 package. Kaplan-Meier analysis was performed in R using the survival package. Accelerated failure time (AFT) regression models were fit using Buckley-James regression with the rms package. Hill coefficients for Dauer and Lifespan data were fit to AFT regression coefficients using the formula E(D) = E0 + (Ef–E0)/(1+(D^–n)*lnIDM)) using braidrm, with E0 fixed and Ef, n, and IDM estimated.

### Data Availability

Strains and reagents are available upon request. The authors affirm that all data necessary for confirming the conclusions are present within the article and its figures. Other relevant data are available within the article and its supplementary information files, or from the corresponding author upon reasonable request.

## Acknowledgements

We thank Joy Alcedo (Wayne State University), Julián Cerón (IDIBELL), Karen Thijssen and Hannes Lans (Erasmus University Medical Center), Rhys McDonough (University of Cambridge), and Matt Ragle and Jordan Ward (University of California, Santa Cruz) for nematode strains; and Christian Frøkjær-Jensen and Sonia El-Mouridi (KAUST) for providing MosTI strains and reagents. We are grateful to the CRG Core Technologies Programme for their support and assistance in this work, including the CRG Advanced Light Microscopy Unit (ALMU). We acknowledge Lucia Sedlackova of the CRG Dynamics of Living Systems Lab for writing the custom Python script for image processing and analysis. This work was technically supported by the EMBL Genomics Core facility. Research for this publication has been partially carried out in the Barcelona Collaboratorium for Modelling and Predictive Biology.

