## Supplementary Figures for "Engineering the auxin-inducible degron system for tunable *in vivo* control of organismal physiology"

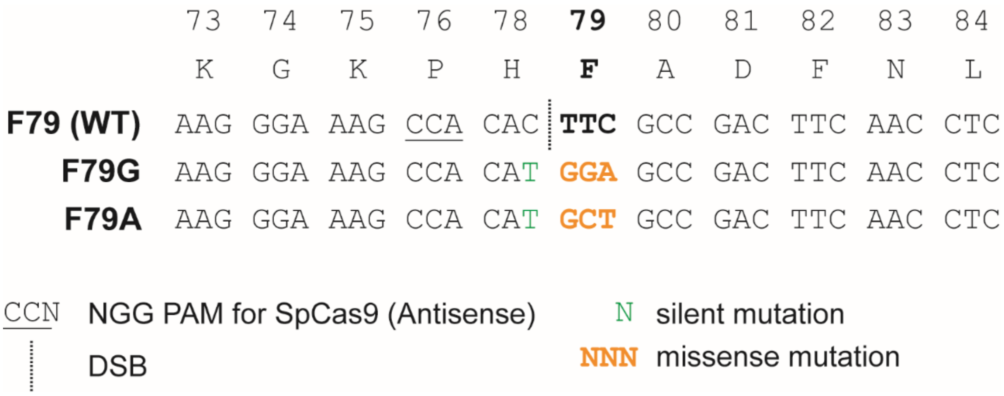


**Supplementary Figure 1**. Conversion of the wild-type *At*TIR1 F79 residue to F79G and F79A using CRISPR–Cas9. SpCas9 directed by a guide RNA (5'–AGGTTGAAGTCGGCGAAGTG–3') produces a double strand break (DSB) immediately upstream of the codon coding for the phenylalanine residue at amino acid position 79. Single-stranded oligodeoxynucleotide (ssODN) donors were used to introduce the corresponding mutations (yellow) that convert phenylalanine into glycine or alanine through homology-directed repair (HDR). Moreover, an additional silent mutation (green) was introduced in amino acid position 78 (c.238C>T) to facilitate genotyping using amplification-refractory mutation system PCR (ARMS-PCR).

### Supplementary Figures

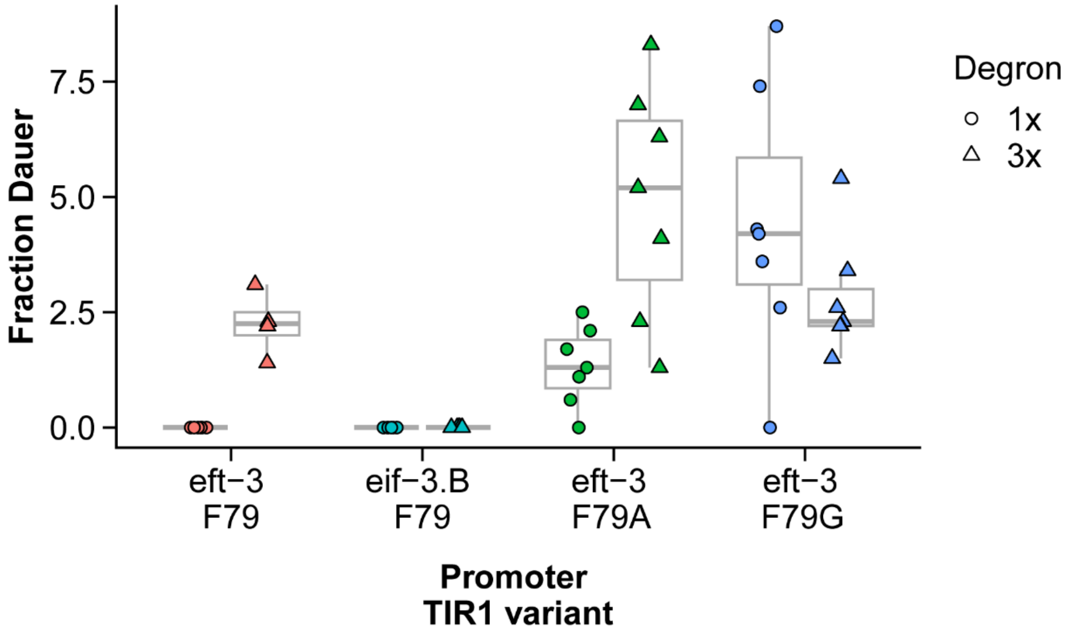


**Supplementary Figure 2.** Auxin-independent effects in different promoter–TIR1 variant combinations in the dauer assay. Even in the absence of auxin, some worms proceed into dauer entry, suggesting the occurrence of DAF-2 basal degradation that results in dauer formation. The weakly expressing *eif-3.Bp* does not result in dauer formation, regardless of degron tag number. In contrast, the strongly expressing *eft-3p* results in dauer formation in all TIR1 variants when there are three degrons, and when there is a single degron in both F79A and F79G, but not in F79. The data is obtained from a single trial with seven replicates each.

### Supplementary Figures

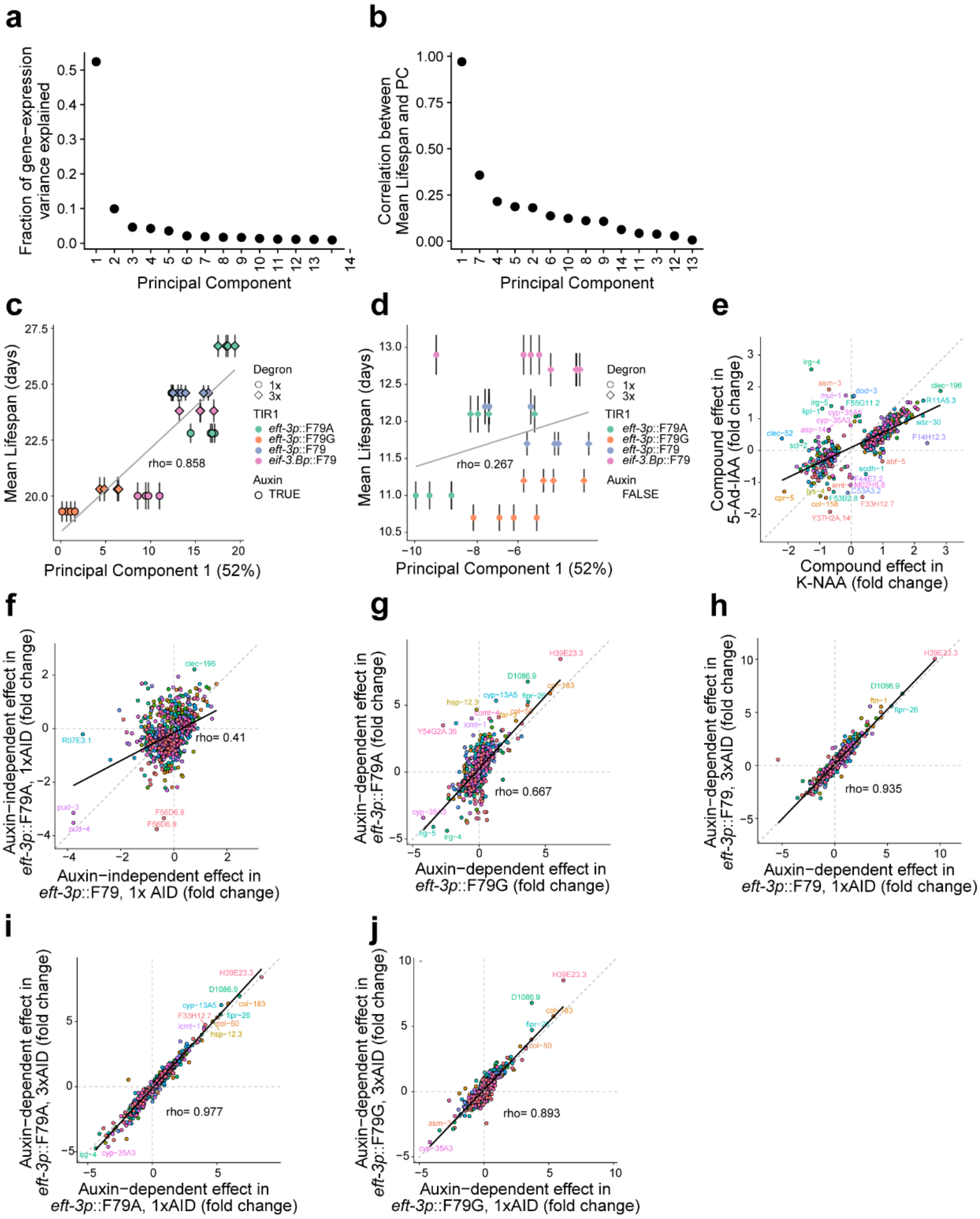


**Supplementary Figure 3.** Correlation of gene expression changes with phenotypic outcomes and between different TIR1–degron pairs. Principal component analysis (PCA) was performed on the transcriptomes of day 1 young adults to determine principal components (PCs) which account for transcriptomic variation. (a) The fraction of gene-expression variance explained by the first 14 PCs across all TIR1 variants. (b) The correlation between each PC and mean lifespan. (c) The correlation between PC1 and mean lifespan considering only auxin-dependent effects or (d) auxin-independent effects. (e) The effect of exposing wild-type animals to K-NAA, compared to the effect of 5-Ad-IAA. (f) In the absence of any compound in DAF-2::AID populations, the effect of *eft-3p*::TIR1[F79] on gene expression compared to the effect of *eft-3p*::TIR1[F79A]. (g) In the presence of their respective activating compounds, the effect of TIR1[F79A] on gene expression compared to the effect of *eft-3p*::TIR1[F79G]; and the effect on gene expression of DAF-2::1xAID vs. DAF-2::3xAID in the (h) *eft-3p*::TIR1[F79], (i) F79A, or (j) F79G backgrounds.

### Supplementary Figures

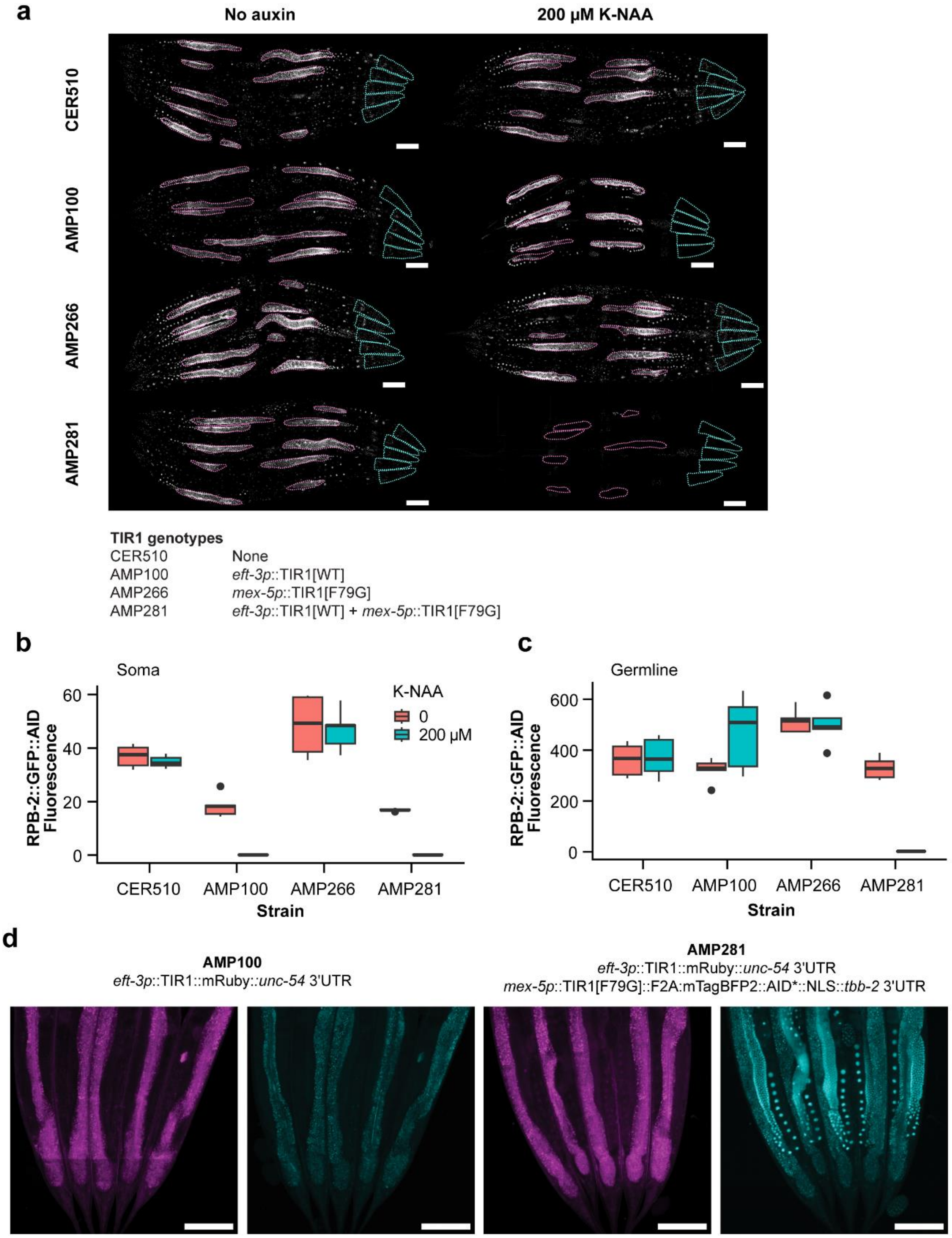


**Supplementary Figure 4**. Trans-activation of *eft-3p*::TIR1::mRuby::*unc-54* 3'UTR in the germline in the presence of *mex-5p*::TIR1[F79G]::*tbb-2* 3'UTR in the dual-channel strain AMP281. (a) Representative images of worms with RPB-2::GFP::degron combined with different TIR1 genotypes without auxin or on 200 µM K-NAA. (b) Quantification of RPB-2::GFP signal in the soma or (c) in the germline of worms shown in (a). The signal is quantified in absolute photon counts from five worms per condition from a single biological replicate. (d) Comparison of *eft-3p*::TIR1::mRuby::*unc-54* 3'UTR signal in the germline in the absence (AMP100) or presence (AMP281) of *mex-5p*::TIR1[F79G]::F2A::mTagBFP2::AID*::NLS:: *tbb-2* 3'UTR. TIR1::mRuby and TIR1[F79G]::mTagBFP2 are imaged in the red channel and blue channels, respectively. Scale bars equal 100 µm.

### Supplementary Figures

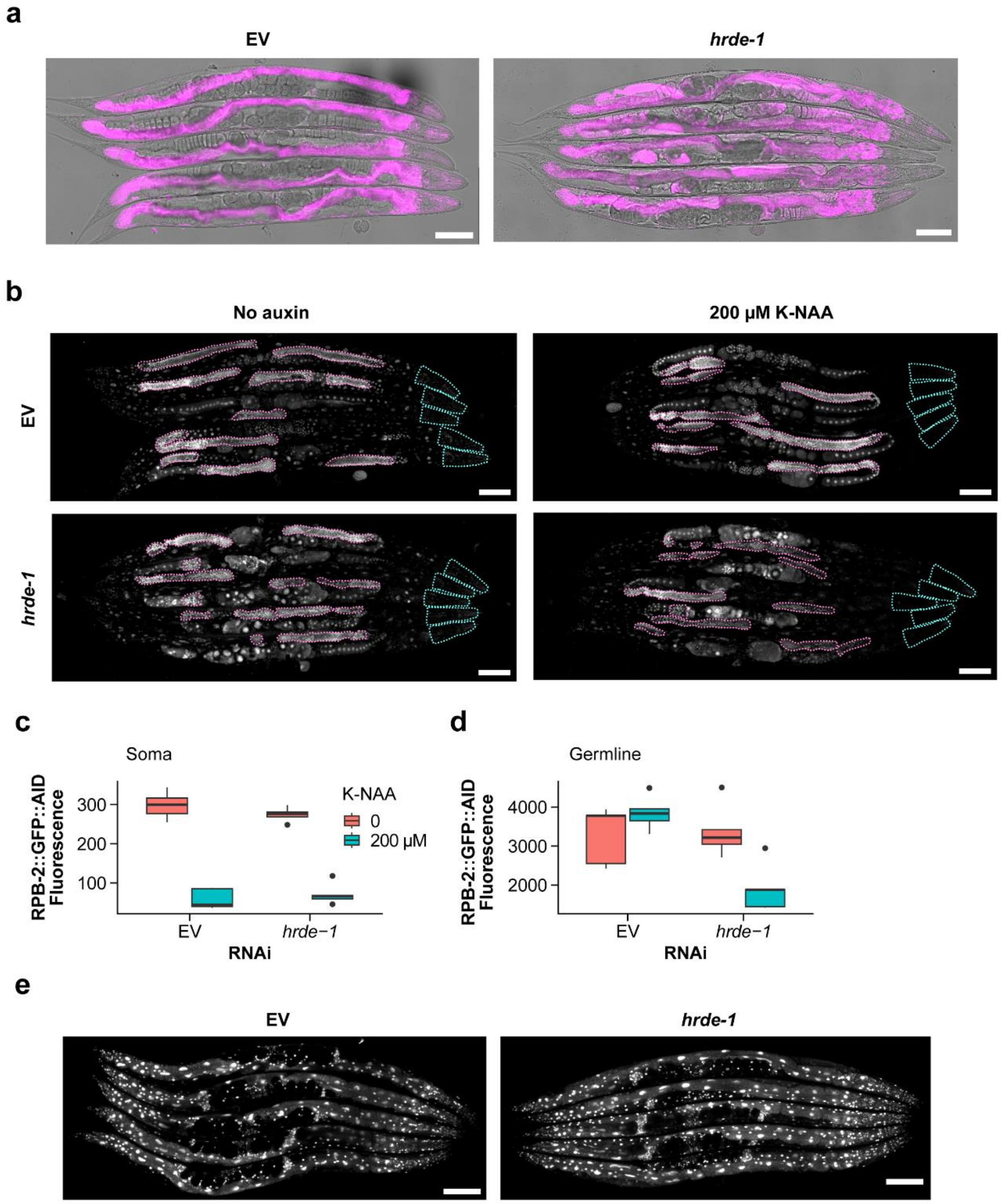


**Supplementary Figure 5**. De-silencing of TIR1 in the germline upon exposure to *hrde-1* RNAi. (a) De-silencing of *eft-3p*::TIR1::mRuby::*unc-54* 3'UTR in the germline of worms exposed to five generations of *hrde-1* RNAi. (b) Depletion of degron-tagged RPB-2::GFP signal in worms with *eft-3p*::TIR1::mRuby::*unc-54* 3'UTR when exposed to five generations of *hrde-1* RNAi and three hours of 200 µM K-NAA. Quantification of RPB-2::GFP signal in the (c) soma (head) or (d) germline of worms shown in (b). The signal is quantified in absolute photon counts from five worms per condition from a single biological replicate. (e) De-silencing of *eft-3p*::TIR1::SL2::NLS::mTagBFP2::*tbb-2* 3'UTR in the germline upon exposure to *hrde-1* RNAi for eight generations. Scale bars equal 100 µm.
