## Supplementary Tables for "Engineering the auxin-inducible degron system for tunable *in vivo* control of organismal physiology"

#### Supplementary Table 1. List of differentially expressed genes in wild-type QZ0 worms upon exposure to auxin analogues

| K-NAA | | | 5-Ad-IAA | 5-Ph-IAA |
|---|---|---|---|---|
| B0281.5 | Y58A7A.5 | *jmjd-3.2* | B0281.5 | F55G11.2 |
| C06E2.5 | Y62H9A.15 | *lbp-7* | C08F1.6 | K02C4.8 |
| C07D10.5 | Y71F9AL.7 | *lec-9* | C17E4.2 | W07E6.5 |
| C08F1.6 | ZK185.9 | *lfor-1* | F33H12.7 | Y37H2A.14 |
| C10C5.4 | ZK550.2 | *lido-8* | F53B2.8 | *adh-1* |
| C10G8.4 | *abts-4* | *mnp-1* | F55G11.2 | *asah-1* |
| C14H10.2 | *acdh-1* | *nhr-2* | K02B12.2 | *asm-3* |
| C17C3.5 | *ags-3* | *nlp-31* | M02H5.8 | *clec-47* |
| C17E4.2 | *aptf-4* | *noah-1* | R11A5.3 | *cyp-35A3* |
| C17E4.20 | *arrd-1* | *oac-32* | T25E12.6 | *cyp-35A5* |
| C31H5.4 | *arrd-2* | *phdh-1* | Y37H2A.14 | *cyp-35C1* |
| C33G3.4 | *asp-5* | *pmp-1* | Y46G5A.7 | *dod-3* |
| C34H4.1 | *atz-1* | *scl-2* | *acdh-1* | *gst-5* |
| C35E7.5 | *btb-11* | *scl-24* | *amt-1* | *hmit-1.1* |
| C39B5.2 | *btb-6* | *sdz-14* | *asah-1* | *irg-4* |
| C50B6.7 | *btb-8* | *sdz-21* | *asm-3* | *irg-5* |
| D1086.7 | *cav-1* | *sdz-28* | *btb-6* | *lfor-1* |
| E02C12.8 | *cec-8* | *sdz-30* | *btb-8* | *lys-4* |
| E04F6.15 | *ceh-39* | *sea-1* | *clec-196* | *nlp-31* |
| F08F1.4 | *cest-13* | *sepa-1* | *clec-47* | *pgph-3* |
| F09F7.6 | *cht-1* | *skr-10* | *col-158* | |
| F14B6.3 | *clc-23* | *skr-13* | *cyp-35A5* | |
| F14H12.3 | *clec-196* | *skr-14* | *cyp-35C1* | |
| F15B10.3 | *clec-265* | *skr-15* | *dod-3* | |
| F15D4.5 | *clec-52* | *skr-7* | *epg-2* | |
| F17C11.6 | *clec-62* | *skr-8* | *fbxb-26* | |
| F22E5.20 | *clec-67* | *skr-9* | *fbxc-36* | |
| F31E8.4 | *cpr-4* | *spp-25* | *fmo-1* | |
| F33E2.5 | *cpr-5* | *sptl-2* | *hmg-11* | |
| F39F10.3 | *cpr-9* | *sup-36* | *inx-2* | |
| F45B8.6 | *cyd-1* | *tatn-1* | *irg-4* | |
| F48C1.9 | *cyn-6* | *tbx-11* | *irg-5* | |
| F48E3.6 | *dhs-9* | *tipn-1* | *lipl-1* | |
| F59A6.12 | *die-1* | *ttr-20* | *lys-4* | |
| H06H21.8 | *drd-5* | *ttr-21* | *mul-1* | |
| H37A05.4 | *dsl-2* | *ttr-50* | *pgph-3* | |
| K02B12.2 | *duxl-1* | *ugt-26* | *pmp-1* | |
| K04G2.10 | *epg-2* | *ule-1* | *sdz-28* | |
| K10D11.5 | *fbxb-10* | *ule-2* | *skr-10* | |
| M02D8.3 | *fbxb-15* | *vet-1* | *skr-7* | |
| R11A5.3 | *fbxb-26* | *vet-2* | *skr-9* | |
| T05D4.2 | *fbxb-91* | *vet-6* | *tatn-1* | |
| T09B4.5 | *fbxc-21* | | *tipn-1* | |
| T25E12.6 | *fbxc-36* | | *vet-6* | |
| T28F3.8 | *fbxc-51* | | | |
| W04A8.4 | *fmo-1* | | | |
| Y106G6D.1 | *gem-4* | | | |
| Y106G6D.2 | *gmap-1* | | | |
| Y15E3A.5 | *gpr-1* | | | |
| Y27F2A.8 | *gst-5* | | | |
| Y32F6A.4 | *hacd-1* | | | |
| Y38H6C.15 | *hil-2* | | | |
| Y45G5AM.5 | *hil-3* | | | |
| Y46G5A.20 | *his-24* | | | |
| Y46G5A.7 | *hmg-11* | | | |
| Y46H3C.5 | *hmit-1.1* | | | |
| Y46H3C.7 | *hsp-16.41* | | | |
| Y51A2D.13 | *inx-2* | | | |

### Supplementary Tables

#### Supplementary Table 2. List of differentially expressed genes in DAF-2::AID and TIR1-expressing strains upon exposure to their corresponding activating ligands

| *eft-3p*::TIR1[F79], *daf-2*::1xAID | | | | *eft-3p*::TIR1[F79], *daf-2*::3xAID | | | | |
|---|---|---|---|---|---|---|---|---|
| B0024.4 | Y51F10.7 | *ech-7* | *srlf-32* | B0024.4 | M02D8.6 | *clec-85* | *gpd-3* | *spi-4* |
| B0513.4 | Y53G8AL.1 | *elo-2* | *tag-244* | C01B10.6 | M28.10 | *clik-3* | *gpdh-1* | *spig-10* |
| C02F12.5 | Y54G2A.36 | *elo-6* | *tbb-6* | C02F12.5 | M60.4 | *col-103* | *grd-3* | *spig-11* |
| C08E8.4 | Y57G11B.5 | *epic-2* | *tbx-43* | C04H5.7 | PDB1.1 | *col-124* | *grd-5* | *spp-1* |
| C09G5.7 | Y69A2AL.2 | *ethe-1* | *test-1* | C07E3.9 | R03H10.6 | *col-129* | *gsnl-1* | *spp-18* |
| C13F10.7 | Y69A2AR.22 | *exc-13* | *ttr-23* | C08E8.4 | R06F6.14 | *col-139* | *gst-20* | *spp-2* |
| C14C6.2 | ZC395.5 | *far-1* | *ttr-26* | C09G5.7 | T01C3.2 | *col-142* | *gst-42* | *sqrd-1* |
| C17F4.7 | ZK354.2 | *far-3* | *ttr-5* | C13F10.7 | T01D1.4 | *col-159* | *gst-5* | *srdh-1* |
| C18A11.1 | ZK380.t2 | *far-8* | *tts-1* | C14C6.5 | T02C5.1 | *col-160* | *hacd-1* | *sri-40* |
| C23H5.8 | ZK863.8 | *fip-6* | *ugt-41* | C17F4.7 | T12D8.5 | *col-183* | *hex-1* | *swt-3* |
| C24B9.3 | *abf-2* | *fipr-26* | *unc-18* | C17H12.6 | T23E7.2 | *col-20* | *hgo-1* | *sym-1* |
| C25E10.8 | *acdh-1* | *folt-2* | | C18A11.1 | W01C8.5 | *col-42* | *his-60* | *taf-11.2* |
| C27C12.1 | *acox-1.4* | *ftn-1* | | C23H5.8 | W01C9.2 | *col-50* | *his-62* | *tag-18* |
| C30G12.2 | *adh-1* | *gadr-6* | | C24B9.3 | W08G11.6 | *col-80* | *hphd-1* | *tag-244* |
| C34H4.2 | *akt-2* | *ges-1* | | C25E10.8 | Y105C5B.15 | *col-81* | *hrg-7* | *tbx-43* |
| C36C9.10 | *aldo-2* | *glb-1* | | C26B9.5 | Y106G6D.2 | *col-93* | *hsp-12.3* | *thn-2* |
| C50B6.7 | *aqp-1* | *gst-20* | | C27A7.6 | Y39B6A.1 | *col-98* | *icl-1* | *tth-1* |
| D1086.3 | *asah-1* | *gst-42* | | C34H4.2 | Y39B6A.5 | *comt-3* | *icmt-1* | *ttr-23* |
| D1086.9 | *asp-3* | *gst-5* | | C36C9.10 | Y43C5A.3 | *comt-4* | *irg-5* | *ttr-26* |
| E01G4.7 | *btb-16* | *hacd-1* | | C49A9.10 | Y51F10.7 | *cpn-3* | *irg-8* | *ttr-5* |
| F01D5.5 | *cdo-1* | *heh-1* | | C50B6.7 | Y53G8AL.1 | *cpn-4* | *ketn-1* | *tts-1* |
| F08F3.4 | *cdr-2* | *his-62* | | C50F7.5 | Y54G2A.36 | *cpr-1* | *klo-1* | *ugt-41* |
| F09E10.15 | *cdr-4* | *hphd-1* | | D1086.3 | Y57G11B.5 | *cpr-4* | *lea-1* | *ugt-63* |
| F13A2.4 | *cest-1.1* | *hrg-7* | | D1086.9 | Y69A2AR.22 | *cpr-5* | *lec-8* | *unc-15* |
| F13D11.4 | *cest-32* | *hsp-12.3* | | E01G4.3 | Y69E1A.5 | *cpr-6* | *lev-11* | *unc-18* |
| F15B9.6 | *citk-1* | *irg-5* | | E01G4.7 | ZK105.1 | *cpr-9* | *lfor-1* | *unc-54* |
| F15E6.3 | *ckr-2* | *irg-8* | | F01D5.1 | ZK1073.1 | *ctsa-1.2* | *lgmn-1* | *unc-87* |
| F19B10.13 | *clc-1* | *lbp-8* | | F01D5.5 | ZK354.2 | *ctsa-2* | *linc-6* | |
| F21C10.11 | *clc-23* | *lea-1* | | F08F3.4 | ZK355.2 | *cyp-13A5* | *linc-7* | |
| F23A7.4 | *clc-24* | *lfor-1* | | F08G2.5 | ZK380.t2 | *cyp-29A2* | *lipl-5* | |
| F23A7.8 | *cld-9* | *linc-6* | | F09C8.1 | ZK637.14 | *cyp-34A9* | *lys-2* | |
| F26D11.1 | *clec-169* | *linc-7* | | F09E10.15 | ZK863.8 | *dct-16* | *lys-4* | |
| F30A10.14 | *clec-52* | *lipl-5* | | F10E9.12 | *abf-2* | *deb-1* | *lys-7* | |
| F30H5.4 | *clec-56* | *lys-2* | | F12A10.1 | *acp-5* | *dgat-2* | *mlc-1* | |
| F35F10.1 | *clec-57* | *lys-4* | | F13A2.4 | *act-4* | *dgk-2* | *mlt-4* | |
| F35F10.5 | *clec-65* | *lys-7* | | F13D11.4 | *adh-1* | *dlg-1* | *mrps-23* | |
| F40F8.4 | *clec-85* | *mlt-4* | | F13H8.11 | *aldo-2* | *dod-23* | *msa-1* | |
| F43C9.2 | *col-129* | *msa-1* | | F15B9.6 | *aptf-4* | *dod-24* | *msra-1* | |
| F47B8.2 | *col-139* | *mtl-1* | | F15E6.3 | *aqp-1* | *dod-3* | *mtl-1* | |
| F48C1.9 | *col-142* | *mtl-2* | | F15G9.1 | *asah-1* | *dod-6* | *mtl-2* | |
| F48D6.4 | *col-159* | *nhr-80* | | F19B10.13 | *asns-2* | *dpy-1* | *myo-3* | |
| F52D2.14 | *col-160* | *nlp-20* | | F21C10.11 | *asp-3* | *dpy-14* | *nlp-20* | |
| F54F7.2 | *col-183* | *nnt-1* | | F23A7.4 | *asp-5* | *drd-50* | *nlp-29* | |
| F55G11.2 | *col-50* | *npa-1* | | F23A7.8 | *asp-6* | *dyf-3* | *nnt-1* | |
| F56D6.8 | *col-80* | *nspc-9* | | F26D11.1 | *bath-12* | *ech-7* | *nspc-10* | |
| F56D6.9 | *col-81* | *pck-1* | | F30H5.4 | *best-14* | *egl-9* | *nspc-13* | |
| F59C12.4 | *col-93* | *pgp-1* | | F33H12.7 | *best-24* | *elo-2* | *nspc-14* | |
| H37A05.4 | *col-96* | *pgp-8* | | F35F10.5 | *bgal-2* | *elo-5* | *nspc-4* | |
| H39E23.3 | *col-98* | *plpp-1.2* | | F42A8.1 | *btb-16* | *elo-6* | *nspc-9* | |
| K09H9.5 | *comt-3* | *poml-3* | | F46F2.3 | *ceeh-1* | *epic-2* | *nspg-10* | |
| K11D12.13 | *comt-4* | *pugs-8* | | F46H5.3 | *cest-1.1* | *ethe-1* | *pck-1* | |
| M60.4 | *cpg-9* | *rimb-1* | | F47B8.2 | *cest-17* | *exc-13* | *pck-2* | |
| PDB1.1 | *cpr-1* | *rpr-1* | | F48C1.9 | *cht-3* | *far-1* | *pcp-3* | |
| R03H10.6 | *cpr-4* | *sepa-1* | | F48D6.4 | *citk-1* | *far-3* | *pdi-2* | |
| T05A12.4 | *cpr-5* | *ser-4* | | F49C12.7 | *ckr-2* | *far-6* | *pgp-8* | |
| T07E3.4 | *cpr-6* | *slc-17.6* | | F52D2.14 | *clc-23* | *far-8* | *plpp-1.2* | |
| T12D8.5 | *cpr-9* | *sod-3* | | F54F7.6 | *clc-24* | *fat-3* | *pmp-5* | |
| T13F3.6 | *cyp-13A5* | *spi-2* | | F55G1.7 | *clc-26* | *fat-5* | *poml-3* | |
| T21C12.8 | *cyp-34A9* | *spig-10* | | F55G11.2 | *cld-9* | *fip-6* | *popl-4* | |
| T28F4.5 | *dct-16* | *spig-9* | | F56D6.9 | *clec-169* | *fipr-26* | *pqn-92* | |
| W04A4.2 | *dod-23* | *spp-1* | | F59C12.4 | *clec-41* | *folt-2* | *pugs-8* | |
| W08G11.6 | *dod-24* | *spp-17* | | F59C6.16 | *clec-52* | *ftn-1* | *rbmx-2* | |
| Y105C5B.15 | *dod-3* | *spp-2* | | H34I24.2 | *clec-56* | *ges-1* | *rpr-1* | |
| Y106G6D.2 | *dod-6* | *srdh-1* | | H39E23.3 | *clec-57* | *glb-1* | *ser-4* | |
| Y17D7C.3 | *dpy-1* | *sri-40* | | K09C4.5 | *clec-65* | *gly-13* | *sld-5* | |
| Y43C5A.3 | *drd-50* | *srlf-30* | | K09H9.5 | *clec-80* | *gpd-2* | *sod-3* | |

### Supplementary Tables

#### *Supplementary Table 2 (continued)*

| *eif-3.Bp*::TIR1[F79], *daf-2*::1xAID | | | | | *eif-3.Bp*::TIR1[F79], *daf-2*::3xAID | | | |
|---|---|---|---|---|---|---|---|---|
| B0281.5 | F53E10.1 | adh-1 | far-5 | pud-4 | C02F12.5 | Y17D7C.3 | cyp-13A5 | pck-2 |
| B0507.3 | F54D11.3 | ahcy-1 | far-8 | pugs-8 | C06B3.6 | Y18D10A.23 | cyp-14A5 | pcp-3 |
| C01B10.3 | F55G1.7 | aly-1 | fat-7 | ras-2 | C07E3.9 | Y43C5A.3 | cyp-34A9 | pgp-8 |
| C02F12.5 | F56D6.8 | aptf-2 | fbf-2 | rimb-1 | C08E8.4 | Y48G1A.2 | cyp-37B1 | pho-14 |
| C04A11.2 | F56D6.9 | aptf-4 | fbl-1 | rml-5 | C09G5.7 | Y49E10.18 | dct-16 | plpp-1.2 |
| C05D9.7 | F59C12.3 | arrd-1 | fbxa-163 | rol-3 | C17C3.5 | Y51F10.7 | del-6 | pmp-5 |
| C07E3.9 | F59C12.4 | atz-1 | fbxb-10 | rpl-11.2 | C17F4.7 | Y53G8AL.1 | dhs-14 | poml-3 |
| C08E8.4 | F59C6.16 | best-14 | fbxb-15 | rpl-23A.1 | C17H12.6 | Y53H1A.2 | dhs-21 | popl-4 |
| C08F1.10 | H11L12.1 | btb-10 | fbxb-26 | rpr-1 | C18A11.1 | Y54G2A.36 | dod-3 | ppw-1 |
| C08F1.6 | H37A05.4 | btb-11 | fbxb-41 | rrf-2 | C18B2.4 | Y58A7A.3 | dpy-1 | pqn-92 |
| C09F9.2 | H39E23.3 | btb-6 | fbxb-91 | rrn-3.56 | C23H5.8 | Y58A7A.5 | ech-7 | pud-3 |
| C09G5.7 | K02B12.2 | catp-3 | fbxc-21 | sac-2 | C24B9.3 | Y69A2AR.22 | elo-2 | pud-4 |
| C13F10.7 | K04G2.10 | cav-1 | fbxc-51 | samt-1 | C34H4.2 | Y94H6A.10 | elo-5 | pugs-8 |
| C14E2.12 | K06A9.1 | cca-1 | fipr-21 | scd-2 | C36C9.10 | ZC395.5 | elo-6 | rml-5 |
| C14H10.2 | K09F6.9 | cdo-1 | fipr-26 | sdc-2 | C49G7.7 | ZK1307.1 | exc-13 | rpl-11.2 |
| C17E4.2 | K10D3.4 | cec-4 | fmi-1 | sdz-14 | C51E3.10 | ZK354.2 | faah-2 | rpr-1 |
| C17E4.20 | K12H4.2 | ceh-20 | frm-8 | sdz-21 | D1086.9 | ZK380.t2 | far-1 | scl-24 |
| C18D4.6 | M01G12.9 | ceh-49 | ftn-1 | sdz-28 | E01G4.7 | ZK863.8 | far-3 | ser-4 |
| C25F9.2 | M03B6.4 | cest-13 | glam-1 | sdz-30 | F08F3.4 | acdh-1 | far-6 | snet-1 |
| C31G12.1 | R02D5.1 | cfh-1 | grd-4 | sepa-1 | F09E10.15 | acox-1.4 | far-8 | sod-3 |
| C35E7.5 | R03H10.6 | chaf-1 | gst-24 | ser-4 | F10E9.12 | adh-1 | fipr-21 | spig-1 |
| C36C9.10 | R04B3.2 | che-3 | haf-9 | sex-1 | F13A2.4 | aldo-2 | fipr-26 | spig-10 |
| C40H1.7 | R05G9.3 | cima-1 | hmg-11 | shc-2 | F13D11.4 | anmt-2 | folt-2 | spig-11 |
| C48B4.10 | R06C1.4 | citk-1 | hrg-10 | sipa-1 | F15B9.6 | aqp-1 | ftn-1 | spig-9 |
| C49G7.12 | R07E3.1 | ckr-2 | hsp-17 | skr-10 | F15E6.3 | aqp-7 | gadr-6 | spp-1 |
| C56E6.2 | R11A5.3 | clec-160 | hum-6 | skr-13 | F19B10.13 | asah-1 | gba-4 | spp-18 |
| D1086.9 | T01D3.1 | clec-196 | icmt-1 | skr-15 | F23A7.4 | asp-3 | ges-1 | spp-2 |
| E01G4.7 | T02D1.8 | clec-47 | irg-5 | skr-16 | F23A7.8 | best-24 | glb-1 | spsb-2 |
| F01D5.5 | T05D4.2 | clec-78 | jmjd-3.2 | skr-7 | F30H5.4 | bgal-2 | glt-1 | srdh-1 |
| F08F3.4 | T06E6.10 | clec-86 | jph-1 | skr-8 | F33H12.7 | btb-16 | gpdh-1 | sri-40 |
| F08G2.5 | T07D3.9 | cls-3 | lea-1 | skr-9 | F35F10.5 | catp-3 | gst-20 | tag-244 |
| F09E10.15 | T08B1.4 | col-129 | lfor-1 | slcr-46.3 | F40F8.4 | cca-1 | gst-24 | tbx-43 |
| F09F7.6 | T18D3.1 | col-139 | lido-11 | sld-5 | F43C9.2 | ceh-20 | gst-5 | thn-2 |
| F10E9.11 | T19D12.6 | col-142 | lido-7 | sod-3 | F46F2.3 | ceh-83 | hacd-1 | timm-17B.2 |
| F10E9.12 | T22B7.4 | col-183 | linc-6 | spi-2 | F47B8.2 | cest-1.1 | his-60 | tni-3 |
| F11D5.1 | T25E12.6 | col-50 | linc-7 | spi-4 | F48C1.9 | cima-1 | his-62 | ttr-23 |
| F12E12.1 | VC5.2 | col-80 | lipl-5 | spig-1 | F48D6.4 | citk-1 | hphd-1 | ttr-26 |
| F13A2.4 | W01C9.2 | col-98 | mak-2 | spig-10 | F49C12.7 | ckr-2 | hrg-7 | ttr-5 |
| F14B6.3 | W05H9.3 | comt-4 | mecr-1 | spig-11 | F52D2.14 | clc-23 | hsp-12.3 | tts-1 |
| F14F9.4 | W08G11.6 | cosa-1 | mes-6 | spig-9 | F55G1.7 | clc-24 | icl-1 | ugt-26 |
| F14H3.3 | W09G12.9 | cpg-9 | mlt-4 | spp-2 | F55G11.2 | cld-9 | icmt-1 | ugt-41 |
| F14H3.4 | Y106G6D.1 | cpn-4 | mnp-1 | spsb-2 | F56D5.6 | clec-160 | irg-3 | ugt-48 |
| F15B9.6 | Y106G6D.2 | cpr-1 | mrp-2 | srdh-1 | F56D6.8 | clec-169 | irg-5 | ugt-62 |
| F15D4.5 | Y11D7A.7 | cpr-4 | mrps-23 | srlf-31 | F56D6.9 | clec-47 | irg-8 | |
| F15E6.3 | Y15E3A.4 | cpr-5 | mth-1 | sup-36 | F59C12.4 | clec-57 | klo-1 | |
| F17A2.13 | Y17D7C.3 | cpr-9 | mtl-1 | taf-11.1 | F59C6.16 | clec-65 | lea-1 | |
| F18E3.11 | Y27F2A.8 | cri-2 | mua-3 | tag-209 | H11L12.1 | col-103 | lfor-1 | |
| F18E3.12 | Y32H12A.8 | crn-6 | mxl-3 | tbx-43 | H34I24.2 | col-129 | lgmn-1 | |
| F18E3.13 | Y39D8A.1 | ctb-1 | nduo-4 | tir-1 | H39E23.3 | col-139 | linc-6 | |
| F18H3.4 | Y45G5AM.5 | cubn-1 | nduo-5 | tni-3 | M01G12.9 | col-142 | linc-7 | |
| F19B10.13 | Y48C3A.18 | cyd-1 | nhr-10 | ttc-36 | PDB1.1 | col-159 | lipl-5 | |
| F21F3.4 | Y49F6C.8 | cyn-10 | nhr-154 | ttr-26 | R03H10.6 | col-183 | lro-1 | |
| F22E5.17 | Y53G8AL.1 | cyp-13A5 | nhr-66 | ttr-50 | R07C12.2 | col-42 | lys-2 | |
| F22E5.20 | Y54G2A.36 | cyp-14A5 | nlp-20 | tts-1 | R11D1.3 | col-50 | lys-4 | |
| F23A7.4 | Y55F3AM.11 | cyp-34A9 | nlp-29 | txdc-12.2 | T02C5.1 | col-80 | lys-7 | |
| F23A7.8 | Y55F3BR.2 | cyp-37B1 | nspc-13 | ugt-26 | T05E12.6 | col-81 | mlt-4 | |
| F26C11.3 | Y56A3A.7 | dao-2 | nspc-14 | ugt-29 | T06E6.10 | col-98 | mrp-2 | |
| F30H5.3 | Y58A7A.3 | dct-17 | nspc-9 | ugt-31 | T12D8.5 | comt-3 | msa-1 | |
| F30H5.4 | Y58A7A.5 | del-6 | oac-32 | ugt-39 | T19D12.6 | comt-4 | mth-1 | |
| F31B12.3 | Y68A4A.13 | dgat-2 | pad-2 | ugt-43 | T21B6.3 | cpg-3 | mtl-1 | |
| F33H2.3 | Y69A2AL.2 | dhc-4 | pgp-5 | ugt-48 | T26H5.9 | cpn-4 | mua-3 | |
| F35F10.5 | Y69A2AR.22 | dmd-9 | pgp-8 | unc-41 | T28F4.5 | cpr-1 | myo-3 | |
| F43B10.1 | ZC513.7 | dpy-1 | phdh-1 | unc-53 | VC5.2 | cpr-3 | nnt-1 | |
| F43C9.2 | ZK105.1 | dyf-3 | ppw-1 | vab-19 | W01C9.2 | cpr-4 | npa-1 | |
| F43G6.10 | ZK354.2 | emre-1 | pqn-92 | zfh-2 | W05H9.3 | cpr-5 | nspc-13 | |
| F48C1.9 | ZK380.t2 | enee-1 | ptr-23 | zig-12 | W08G11.6 | cpr-6 | nspc-4 | |
| F48E3.6 | ZK863.8 | epg-2 | ptr-3 | zip-10 | Y105C5B.15 | cpr-9 | oac-32 | |
| | acy-1 | far-3 | pud-3 | ztf-16 | Y11D7A.3 | cri-2 | papl-1 | |

### Supplementary Tables

#### *Supplementary Table 2 (continued)*

| *eft-3p*::TIR1[F79A], *daf-2*::1xAID | | | | | *eft-3p*::TIR1[F79A], *daf-2*::3xAID | | | |
|---|---|---|---|---|---|---|---|---|
| C02F12.5 | R03H10.1 | *col-160* | *lbp-7* | *ugt-26* | B0250.7 | F53A9.8 | ZK228.4 | *cyp-34A9* |
| C07E3.9 | R03H10.6 | *col-183* | *lea-1* | *ugt-31* | B0303.11 | F53B2.8 | ZK354.2 | *cyp-35A2* |
| C08E8.4 | R06C1.4 | *col-50* | *lfor-1* | *ugt-41* | C02F12.5 | F54D11.3 | ZK380.t2 | *cyp-35A3* |
| C10C5.4 | R102.4 | *col-80* | *lgmn-1* | *ugt-46* | C07E3.9 | F54F7.3 | ZK863.8 | *cyp-35C1* |
| C14C6.2 | T02D1.8 | *col-81* | *linc-6* | *ugt-48* | C08E8.4 | F55B11.2 | *abf-2* | *dct-8* |
| C14C6.5 | T12D8.5 | *col-98* | *linc-7* | *ugt-62* | C10C5.4 | F55G1.7 | *abf-5* | *dod-19* |
| C17C3.5 | T13F3.6 | *comt-3* | *lips-10* | *ugt-63* | C14C6.2 | F55G11.2 | *abf-6* | *dod-23* |
| C18A11.1 | T16G12.1 | *comt-4* | *lro-1* | *ugt-8* | C14C6.5 | F55G11.4 | *acp-6* | *dod-3* |
| C18B2.4 | T19B10.2 | *cpg-7* | *lys-4* | *ule-2* | C14E2.12 | F56D6.8 | *acs-17* | *dod-6* |
| C25E10.8 | T21B6.3 | *cpg-9* | *lys-7* | *unc-18* | C17C3.5 | F56D6.9 | *adh-1* | *dpy-1* |
| C27H5.2 | T28F4.5 | *cpr-1* | *mct-3* | *vhp-1* | C17H12.6 | F59C12.4 | *akt-2* | *dpy-14* |
| C31H5.6 | W01C9.2 | *cpr-3* | *mlt-4* | *vit-1* | C18A11.1 | F59C6.16 | *aldo-2* | *drd-1* |
| C36C9.10 | W02B12.1 | *cpr-4* | *msa-1* | *vit-5* | C18B2.4 | H06H21.8 | *amt-4* | *drd-5* |
| C49G7.7 | W05H9.3 | *cpr-5* | *mtl-1* | *xbp-1* | C18E9.5 | H39E23.3 | *anmt-2* | *drd-50* |
| C50F7.5 | W08G11.6 | *ctsa-4.2* | *mtl-2* | *xdh-1* | C25E10.10 | K07C11.7 | *anp-1* | *ech-7* |
| C51E3.10 | W09G12.9 | *cyp-13A5* | *mua-3* | *yap-1* | C25E10.8 | K08C7.6 | *asm-3* | *elo-5* |
| C53B7.2 | Y110A2AL.9 | *cyp-33C8* | *myo-3* | *zig-7* | C27H5.2 | K08D8.4 | *asp-14* | *enpl-1* |
| C53B7.3 | Y20F4.4 | *cyp-34A9* | *nhx-2* | | C30G12.2 | K09E2.3 | *bath-12* | *exc-13* |
| D1086.9 | Y37H2A.14 | *cyp-35A2* | *nnt-1* | | C31H5.6 | K09H9.5 | *btb-14* | *far-1* |
| E01G4.5 | Y38E10A.14 | *cyp-35A3* | *nrf-6* | | C32H11.4 | K11D12.13 | *cah-5* | *far-3* |
| F01D5.1 | Y39B6A.1 | *cyp-35C1* | *oac-14* | | C34H4.2 | M01G12.9 | *catp-7* | *far-5* |
| F01D5.5 | Y43C5A.3 | *dct-8* | *oac-32* | | C36C9.10 | M01H9.3 | *cbl-1* | *far-6* |
| F08F3.4 | Y46G5A.20 | *dhs-21* | *pck-1* | | C41G11.1 | M02D8.6 | *cca-1* | *fat-7* |
| F09E10.15 | Y47G6A.33 | *dod-23* | *pcp-2* | | C49G7.7 | M02H5.8 | *cdr-2* | *fbxa-163* |
| F10E9.12 | Y47H10A.5 | *dod-24* | *pdi-2* | | C50F7.5 | PDB1.1 | *cdr-4* | *fip-6* |
| F12A10.1 | Y51F10.7 | *dod-3* | *pdi-6* | | C51E3.10 | R03H10.1 | *ceh-99* | *fipr-21* |
| F13D11.4 | Y53G8AL.1 | *dod-6* | *pgp-5* | | C53B7.2 | R03H10.6 | *cest-1.1* | *fipr-26* |
| F14F9.4 | Y54G2A.36 | *dpy-1* | *pgp-8* | | C53B7.3 | R102.4 | *cest-13* | *frm-8* |
| F15B9.6 | Y58A7A.5 | *drd-1* | *pgph-3* | | D1086.9 | T01D1.4 | *cest-17* | *ftn-1* |
| F18E3.11 | Y69H2.3 | *drd-5* | *plpp-* | | E01G4.7 | T03D8.6 | *cht-3* | *gadr-6* |
| F18E3.13 | Y94H6A.10 | *drd-50* | *1.2* | | E04F6.15 | T05A12.4 | *ciao-1* | *ges-1* |
| F19B10.13 | ZK228.4 | *dyf-3* | *pnk-1* | | E04F6.9 | T05E12.3 | *citk-1* | *glb-1* |
| F19B2.5 | ZK354.2 | *ech-7* | *poml-3* | | F01D5.1 | T07D3.9 | *ckr-2* | *gldc-1* |
| F21C10.11 | ZK355.2 | *elo-5* | *ppat-1* | | F01D5.5 | T08B1.1 | *clc-1* | *gnrr-2* |
| F23A7.4 | ZK380.t2 | *elo-6* | *pud-3* | | F08F3.4 | T12D8.5 | *clc-24* | *grd-4* |
| F23A7.8 | ZK863.8 | *enpl-1* | *pud-4* | | F09E10.15 | T13F3.6 | *cld-9* | *gst-10* |
| F28H7.3 | *abf-2* | *exc-13* | *pugs-8* | | F10E9.12 | T16G12.1 | *clec-169* | *gst-24* |
| F30H5.4 | *abf-5* | *faah-2* | *rpl-11.2* | | F12A10.1 | T19B10.2 | *clec-41* | *gst-5* |
| F32A5.4 | *adh-1* | *famk-1* | *scl-2* | | F13A2.4 | T21B6.3 | *clec-47* | *gst-6* |
| F33H12.7 | *akt-2* | *far-1* | *scl-24* | | F13D11.4 | T23B3.2 | *clec-48* | *hacd-1* |
| F35B12.9 | *aldo-2* | *far-3* | *ser-4* | | F13D12.3 | T28F4.5 | *clec-49* | *haf-9* |
| F35D11.4 | *amt-4* | *fat-7* | *skr-4* | | F13H8.11 | W01C9.2 | *clec-5* | *hil-5* |
| F35F10.5 | *anp-1* | *fbxa-107* | *slc-17.6* | | F14F9.4 | W02B12.1 | *clec-51* | *him-4* |
| F46F2.3 | *asm-3* | *fbxa-163* | *sod-3* | | F14H12.3 | W05H9.3 | *clec-52* | *his-60* |
| F47B8.2 | *bath-12* | *fip-6* | *spi-2* | | F15B9.6 | W08G11.6 | *clec-57* | *hpo-6* |
| F48C1.9 | *bcmo-2* | *fipr-26* | *spi-4* | | F15E6.3 | W09G12.9 | *cnc-4* | *hsp-12.3* |
| F48D6.4 | *btb-14* | *frm-8* | *spp-18* | | F16B4.4 | Y16B4A.2 | *col-139* | *hsp-16.2* |
| F52D2.14 | *cdo-1* | *ftn-1* | *spp-2* | | F18C5.10 | Y22D7AL.15 | *col-142* | *hsp-16.41* |
| F53B2.8 | *cdr-2* | *gadr-6* | *spp-4* | | F18E3.13 | Y32F6B.1 | *col-159* | *hsp-3* |
| F54F7.3 | *cdr-4* | *ges-1* | *sqst-1* | | F19B10.13 | Y37H2A.14 | *col-183* | *icmt-1* |
| F55B11.2 | *ceh-99* | *glb-1* | *srdh-1* | | F19B2.5 | Y38E10A.14 | *col-19* | *idpp-16* |
| F55G1.7 | *cest-1.1* | *gnrr-2* | *sri-40* | | F21C10.11 | Y39B6A.1 | *col-50* | *ifb-2* |
| F55G11.2 | *cht-3* | *gpd-2* | *srlf-30* | | F23A7.4 | Y39B6A.5 | *col-80* | *ilys-5* |
| F55G11.4 | *citk-1* | *grd-4* | *srlf-35* | | F23A7.8 | Y41E3.8 | *col-81* | *inx-15* |
| F56D6.8 | *ckr-2* | *hacd-1* | *swt-3* | | F28B4.3 | Y43C5A.3 | *comt-3* | *irg-3* |
| F56D6.9 | *clc-24* | *haf-9* | *swt-6* | | F28H7.3 | Y46G5A.20 | *comt-4* | *irg-4* |
| F59C12.4 | *cld-9* | *him-4* | *tag-120* | | F30H5.4 | Y47G6A.33 | *cpg-3* | *irg-5* |
| F59C6.16 | *clec-166* | *his-60* | *tbx-43* | | F32A5.4 | Y47H10A.5 | *cpg-7* | *klo-1* |
| H39E23.3 | *clec-169* | *his-64* | *test-1* | | F33H12.7 | Y48G1A.2 | *cpg-9* | *kynu-1* |
| K08D8.4 | *clec-47* | *hpo-6* | *thn-2* | | F35B12.9 | Y51F10.7 | *cpna-2* | *lbp-7* |
| K09C4.5 | *clec-49* | *hrg-7* | *ttr-23* | | F35D11.4 | Y53G8AL.1 | *cpr-1* | *lea-1* |
| K09E2.3 | *clec-51* | *hsp-12.3* | *ttr-26* | | F35F10.5 | Y54G2A.36 | *cpr-3* | *lfor-1* |
| K09H9.5 | *clec-56* | *icmt-1* | *ttr-29* | | F45D3.4 | Y54G2A.45 | *cpr-5* | *lgmn-1* |
| M01G12.9 | *clec-57* | *idpp-16* | *ttr-42* | | F47B8.2 | Y58A7A.3 | *crt-1* | *linc-6* |
| M01H9.3 | *cnc-4* | *irg-3* | *ttr-5* | | F48C1.9 | Y69H2.3 | *ctsa-3.1* | *linc-7* |
| M02D8.6 | *col-139* | *irg-4* | *tts-1* | | F48D6.4 | Y94H6A.10 | *ctsa-4.2* | *lro-1* |
| M02H5.8 | *col-142* | *irg-5* | *txt-17* | | F48E3.6 | ZC443.3 | *cyp-13A5* | *lys-4* |
| PDB1.1 | *col-159* | *klo-1* | *ugt-22* | | F52D2.14 | ZK1055.7 | *cyp-33C8* | *lys-7* |

### Supplementary Tables

#### *Supplementary Table 2 (continued)*

| *eft-3p*::TIR1[F79A], *daf-2*::3xAID | | *eft-3p*::TIR1[F79G], *daf-2*::1xAID | | *eft-3p*::TIR1[F79G], *daf-2*::3xAID | |
|---|---|---|---|---|---|
| *mlt-4* | *tli-1* | AC3.5 | *dod-3* | C02F12.5 | *clec-169* |
| *msa-1* | *tni-3* | C08F1.6 | *dyf-3* | C08E8.4 | *clec-41* |
| *mtl-1* | *tth-1* | C09F9.2 | *egl-15* | C10C5.4 | *clec-57* |
| *mtl-2* | *ttr-23* | C14E2.12 | *endu-2* | C14C6.2 | *clec-85* |
| *mua-3* | *ttr-26* | C17H12.8 | *faah-2* | C14C6.5 | *col-139* |
| *nape-2* | *ttr-29* | C25E10.8 | *far-3* | C17H12.8 | *col-142* |
| *ndg-4* | *ttr-33* | C30G12.2 | *fat-7* | C18A11.1 | *col-183* |
| *nhr-76* | *ttr-42* | C32E12.4 | *fbxb-91* | C25E10.10 | *col-19* |
| *nhx-2* | *ttr-5* | C40H1.7 | *fbxc-18* | C25E10.8 | *col-50* |
| *nnt-1* | *tts-1* | C53B7.3 | *fipr-26* | C36C9.10 | *col-80* |
| *nrf-6* | *ugt-22* | D1086.9 | *grd-4* | C40H1.7 | *comt-3* |
| *oac-14* | *ugt-26* | E04F6.15 | *gst-10* | C53B7.3 | *cpr-1* |
| *oac-20* | *ugt-31* | F09F7.6 | *hil-3* | D1086.9 | *cpr-4* |
| *oac-32* | *ugt-41* | F12A10.1 | *his-64* | E01G4.5 | *cpr-5* |
| *orai-1* | *ugt-46* | F14B6.3 | *ifb-2* | F01D5.1 | *ctsa-4.2* |
| *pat-2* | *ugt-48* | F14F9.4 | *irg-4* | F01D5.3 | *cyp-13A5* |
| *pck-1* | *ugt-63* | F26C11.3 | *irg-5* | F01D5.5 | *cyp-35A2* |
| *pcp-1* | *ugt-8* | F32A5.4 | *lam-3* | F12A10.1 | *cyp-35A3* |
| *pcp-2* | *ule-2* | F32D8.12 | *lbp-7* | F14F9.4 | *cyp-35C1* |
| *pcp-3* | *unc-18* | F33H12.7 | *lys-4* | F14H12.3 | *dct-8* |
| *pdi-2* | *unk-1* | F48C1.9 | *msa-1* | F15B9.6 | *dod-23* |
| *pdi-6* | *vhp-1* | F48F7.6 | *mua-3* | F15E6.3 | *dod-6* |
| *pept-1* | *vit-1* | F55G11.2 | *nas-11* | F19B10.13 | *dpf-2* |
| *pgp-5* | *vit-3* | F55G11.4 | *nhr-88* | F19B2.5 | *drd-5* |
| *pgp-8* | *vit-4* | F56D6.8 | *noah-1* | F21C10.10 | *far-3* |
| *pho-11* | *vit-5* | F56D6.9 | *pgp-8* | F21C10.11 | *far-5* |
| *plpp-1.2* | *zig-12* | H39E23.3 | *pix-1* | F30A10.14 | *fipr-21* |
| *pnk-1* | *zig-7* | K11D12.13 | *pugs-8* | F32A5.4 | *fipr-26* |
| *poml-3* | | M02H5.8 | *rpl-11.2* | F33H12.7 | *ges-1* |
| *ppat-1* | | MTCE.33 | *rrn-1.2* | F35F10.5 | *glb-1* |
| *pud-3* | | R06C1.4 | *scb-1* | F36A2.3 | *grd-4* |
| *pud-4* | | R07B7.10 | *scl-2* | F46F2.3 | *his-64* |
| *pugs-8* | | T03D8.6 | *scl-24* | F47B8.2 | *hmit-1.1* |
| *rimb-1* | | T05D4.2 | *scl-5* | F48C1.9 | *hpo-6* |
| *rpl-11.2* | | T22B7.4 | *sdc-3* | F53A9.8 | *hrg-7* |
| *rpr-1* | | Y37H2A.14 | *spi-2* | F54F7.3 | *irg-4* |
| *sams-1* | | Y38E10A.14 | *srdh-1* | F55G11.4 | *irg-5* |
| *scl-2* | | Y42G9A.3 | *srlf-30* | F56D6.8 | *lys-4* |
| *ser-4* | | Y43C5A.3 | *srlf-35* | F56D6.9 | *lys-7* |
| *skpo-3* | | Y54G2A.36 | *tag-273* | F59A6.12 | *mlt-7* |
| *skr-4* | | Y58A7A.3 | *tni-3* | H39E23.3 | *msa-1* |
| *slc-17.6* | | Y58A7A.5 | *ttr-26* | K02F3.9 | *mtl-1* |
| *smim-14* | | ZK355.2 | *ttr-29* | K08D8.4 | *mtl-2* |
| *sod-3* | | ZK863.8 | *tub-2* | K09C4.5 | *pgp-8* |
| *spe-15* | | *abf-2* | *ugt-22* | K11D12.13 | *pud-4* |
| *spi-2* | | *acs-7* | *ugt-26* | M02H5.8 | *pugs-8* |
| *spi-4* | | *adh-1* | *ugt-31* | M04D5.3 | *scl-2* |
| *spp-18* | | *asm-3* | *ugt-41* | T05E12.6 | *sod-3* |
| *spp-2* | | *btb-11* | *ugt-48* | T12D8.5 | *spi-2* |
| *spp-4* | | *ceh-20* | *unc-18* | T13F3.6 | *spp-2* |
| *sqst-1* | | *cest-33* | | T19B10.2 | *srdh-1* |
| *srdh-1* | | *citk-1* | | Y105C5B.15 | *sri-40* |
| *sri-40* | | *clec-52* | | Y37H2A.14 | *srlf-30* |
| *srlf-30* | | *clec-57* | | Y38E10A.14 | *srlf-35* |
| *srlf-35* | | *clec-78* | | Y43C5A.3 | *swt-6* |
| *stdh-1* | | *col-183* | | Y58A7A.3 | *tag-244* |
| *sto-1* | | *col-19* | | Y58A7A.5 | *ttr-26* |
| *swt-3* | | *col-50* | | ZK355.2 | *ttr-29* |
| *swt-6* | | *col-93* | | ZK863.8 | *ugt-22* |
| *swt-7* | | *cpn-4* | | *abf-2* | *ugt-26* |
| *tag-120* | | *cpna-2* | | *adh-1* | *ugt-41* |
| *tag-244* | | *cpr-4* | | *asm-3* | *ugt-44* |
| *tbx-43* | | *cpr-5* | | *cdr-2* | *ule-2* |
| *test-1* | | *crn-6* | | *ceh-20* | *unc-18* |
| *thn-2* | | *cyp-35A2* | | *citk-1* | *xdh-1* |
| | | *cyp-35A3* | | *ckr-2* | |
| | | *cyp-35C1* | | *clc-24* | |
| | | *dct-8* | | *clec-166* | |

### Supplementary Tables

#### Supplementary Table 3. List of Strains Used in this Study

| Strain Name | Genotype | Source |
|---|---|---|
| AMP100 | *ieSi57 [eft-3p::TIR1::mRuby::unc-54 3'UTR] II; rpb-2(cer135[rpb-2::GFP*$^{\Delta piRNA}$*::AID::3xFLAG]) III* | Natasha Oswal et al., 2022 |
| AMP145 | *ieSi57 [eft-3p::TIR1::mRuby::unc-54 3'UTR + Cbr-unc-119(+)] II; daf-2(ohm13)[daf-2::AID::3xFLAG] III* | This study |
| AMP158 | *ohm10[eft-3p::TIR1[F79A]::mRuby::unc-54 3'UTR + Cbr-unc-119(+)] II* | This study |
| AMP167 | *ieSi57 [eft-3p::TIR1::mRuby::unc-54 3'UTR + Cbr-unc-119(+)] II; daf-2(ohm17)[daf-2::3xAID::3xFLAG] III* | This study |
| AMP169 | *weSi174[eif-3.Bp::TIR1::linker::mCherry*$^{\Delta piRNA}$*::tbb-2 3'UTR; unc-119(+)] II; daf-2(ohm17)[daf-2::3xAID::3xFLAG] III* | This study |
| AMP175 | *ohm24(unc-119p::TIR1[F79G]::mRuby) IV* | This study |
| AMP184 | *ohm8[eft-3p::TIR1[F79G]::mRuby::unc-54 3'UTR + Cbr-unc-119(+)] II* | This study |
| AMP205 | *weSi174[eif-3.Bp::TIR1::linker::mCherry*$^{\Delta piRNA}$*::tbb-2 3'UTR; unc-119(+)] II; daf-2(ohm13)[daf-2::AID::3xFLAG] III* | This study |
| AMP206 | *ieSi61 [ges-1p::TIR1::mRuby::unc-54 3'UTR + Cbr-unc-119(+)] II; ohm24(unc-119p::TIR1[F79G]::mRuby) IV; hcf-1(cer159[hcf-1::GFP*$^{\Delta piRNA}$*::degron::3xFLAG]) IV* | This study |
| AMP207 | *ohm8[eft-3p::TIR1[F79G]::mRuby::unc-54 3'UTR + Cbr-unc-119(+)] II; daf-2(ohm17)[daf-2::3xAID::3xFLAG] III* | This study |
| AMP208 | *ohm10[eft-3p::TIR1[F79A]::mRuby::unc-54 3'UTR + Cbr-unc-119(+)] II; daf-2(ohm17)[daf-2::3xAID::3xFLAG] III* | This study |
| AMP216 | *ohm8[eft-3p::TIR1[F79G]::mRuby::unc-54 3'UTR + Cbr-unc-119(+)] II; daf-2(ohm13)[daf-2::AID::3xFLAG] III* | This study |
| AMP217 | *ohm10[eft-3p::TIR1[F79A]::mRuby::unc-54 3'UTR + Cbr-unc-119(+)] II; daf-2(ohm13)[daf-2::AID::3xFLAG] III* | This study |
| AMP245 | *ohm49[eft-3p::nTIR1::SL2::NLS::mTagBFP2::tbb-2 3'UTR] IV* | This study |
| AMP266 | *ohm52 [mex-5p::TIR1[F79G]::F2A::mTagBFP2::AID*::NLS::tbb-2 3'UTR] I rpb-2(cer135[rpb-2::GFP*$^{\Delta piRNA}$*::AID::3xFLAG]) III* | This study |
| AMP267 | *ohm53[eft-3p::nTIR1::SL2::NLS::mTagBFP2::eft-3 3'UTR] IV* | This study |
| AMP281 | *ohm52 [mex-5p::TIR1[F79G]::F2A::mTagBFP2::AID*::NLS::tbb-2 3'UTR] I; ieSi57 [eft-3p::TIR1::mRuby::unc-54 3'UTR + Cbr-unc-119(+)] II; rpb-2(cer135[rpb-2:: GFP*$^{\Delta piRNA}$*::AID::3xFLAG]) III* | This study |
| AMP284 | *rpb-2(cer135[rpb-2::GFP*$^{\Delta piRNA}$*::AID::3xFLAG]) III; ohm53[eft-3p::nTIR1::SL2::NLS::mTagBFP2::eft-3 3'UTR] IV* | This study |
| CA1200 | *ieSi57 [eft-3p::TIR1::mRuby::unc-54 3'UTR + Cbr-unc-119(+)] II; unc-119(ed3) III* | Abby Dernburg (CGC) |
| CA1209 | *ieSi61[ges-1p::TIR1::mRuby::unc-54 3'UTR + Cbr-unc-119(+)] II; unc-119(ed3) III* | Abby Dernburg (CGC) |
| CER510 | *rpb-2(cer135[rpb-2::GFP*$^{\Delta piRNA}$*::AID::3xFLAG]) III* | Julián Cerón |
| CER556 | *hcf-1(cer159[hcf-1::GFP*$^{\Delta piRNA}$*::AID::3xFLAG]) IV* | Julián Cerón |
| CFJ94 | *unc-119(ed3) III; kstSi37 [Cbr-unc-119(kst13)] IV* | Christian Frøkjær-Jensen (CGC) |
| EG6699 | *ttTi5605 II; unc-119(ed3) III; oxEx1578* | Christian Frøkjær-Jensen (CGC) |
| HAL227 | *unc-119(ed3) III; emcSi70 [unc-119p::TIR1::mRuby] IV* | Hannes Lans (CGC) |
| JA1880 | *weSi174[eif-3.Bp::TIR1::linker::mCherry*$^{\Delta piRNA}$*::tbb-2 3'UTR; unc-119(+)] II; unc-119(ed3) III* | Rhys McDonough and Julie Ahringer |
| JDW221 | *wrdSi50 [mex-5p::TIR1::F2A::mTagBFP2::AID*::NLS::tbb-2 3'UTR] I* | Jordan Ward (CGC) |
| N2 | Wild type (Bristol N2) | CGC |
| QZ0 | Wild type (Bristol N2) | Joy Alcedo |

### Supplementary Tables

#### Supplementary Table 4. List of Oligos and Constructs

##### *Primers*

| Gene | Forward primer (5'–3') | Reverse primer (5'–3') |
|---|---|---|
| *daf-2* C-terminus | TTTCGGTGAAAATGAGCATCTA | ACGGGAAGTTTTTGATGGTTTT |
| TIR1 External (Chr II) | TGGAAATGCTCGGAAGGACT | TGATGTCCGATTGCAGCTTG |
| TIR1 External (Chr IV) | GGTCCCCATTTCACCAGAGA | GTGGAGGGGCAGTACAGAAT |
| TIR1[F79]-specific | TCAAGGGAAAGCCACACTTC | N/A |
| TIR1[F79G]-specific | TCAAGGGAAAGCCACATGGA | N/A |
| TIR1[F79A]-specific | TCAAGGGAAAGCCACATGCT | N/A |
| TIR1 Internal | N/A | GAAGTGGGAGAGCCAGTGTC |
| *eif-3.Bp* genotyping primers | CGCGGCCTAGGATTCTCTTC (on *eif-3.Bp*) | GAGGAGAGGACGAGGACCTT (on TIR1) |
| *eft-3* promoter | CAACTTCCATTGGTTCTTCCATTGTTTCTG | GGCTGCTACGGAGTGAGCAA |
| *eft-3* 3'UTR swap | AGGTAGCTGTAGCGCGATAT | TAAGACATTGCCGCACAGATT |
| *rpb-2* C-terminus | GTAAGCTGCTCTTCCAGGAGT | TTAACCGGAAAAGTCCGTGAT |
| *hcf-1* C-terminus | ATATGGCCCGGCTACTCAAG | GCGGCAAAGTTGGAAAAGGT |

##### *crRNAs*

| Description | Sequence (5'–3', without PAM) |
|---|---|
| AtTIR1[F79] to AtTIR1[F79A/G] | AGGTTGAAGTCGGCGAAGTG |
| *daf-2* C-terminus | TTTTGGGGGTTTCAGACAAG |
| nTIR1 *tbb-2* to *eft-3* 3'UTR swap N-terminus | AACTTGTGTCCCAGTTTCGA |
| nTIR1 *tbb-2* to *eft-3* 3'UTR swap C-terminus | AAAGTCAGGTCTCTGAGCTC |

##### *ssDNA Repair templates*

| Description | Sequence |
|---|---|
| AtTIR1[F79] to AtTIR1[F79G] | AAGGTCCGTTCCGTCGAGCTCAAGGGAAAGCCACATGGAGCCGACTTCAACCTCGTCCCAGAC GGATGGGGAGG |
| AtTIR1[F79] to AtTIR1[F79A] | AAGGTCCGTTCCGTCGAGCTCAAGGGAAAGCCACATGCTGCCGACTTCAACCTCGTCCCAGAC GGATGGGGAGG |
| nTIR1 *tbb-2* to *eft-3* 3'UTR swap | GTAGCTGTAGCGCGATATTGCGATTTGCCATCAAAGCTTGGACATAAACTTAATTAAATCTTCATTG TTGAGTTTATCTTGTTGATTTTTGAATAAATTATCAACTCTTTACTTTTTAATGGGTTATGAAATAAATAA ACATTGAAAACTGATAAACAACGTTCATCTCTCTCAGGAAACGGAGAATCTGTGCGGCAATG |

##### *Constructs*

| Name | Description | Source |
|---|---|---|
| **pSEM246** | mosTI(*unc-119*) MCS cloning vector (AmpR) | Christian Frøkjær-Jensen (Addgene plasmid # 159821) |
| **pNES0036** | MosTI nTIR1 *unc-119* targeting vector (pSEM246 backbone) | This study |

### Supplementary Tables

### Supplementary Table 5. AID and nTIR1 sequences

#### >1xAID nucleotide sequence

GGATCCGGAGGAGGAGGACCCAAAGGACCCAGCCAAGCCACCAGCCAAGGCCCAAGTCGTCGGATGGCCACCAGTCCGTTCCTACC
GTAAGAACGTCATGGTCTCCTGCCAAAAGTCCTCCGGAGGACCAGAGGCCGCCGCCTTCGTCAAGGAGAACCTCTACTTCCAATCCGGA
AAGGACTACAAGGACCACGACGGAGACTACAAGGACCACGACATCGACTACAAGGACGACGACGACAAG

#### >1xAID amino acid sequence

GSGGGGPKDPAKPPAKAQVVGWPPVRSYRKNVMVSCQKSSGGPEAAAFVKENLYFQSGKDYKDHDGDYKDHDIDYKDDDDK

#### >3xAID nucleotide sequence

GGATCCGGAGGAGGAGGACCCAAAGGACCCAGCCAAGCCACCAGCCAAGGCCCAAGTCGTCGGATGGCCACCAGTCCGTTCCTACC
GTAAGAACGTCATGGTCTCCTGCCAAAAGTCCTCCGGAGGACCAGAGGCCGCCGCCTTCGTCAAGGGAGCTGGAGCCGGAGCTAAGG
AGCCAAAGGATCCAGCTAAGCCACCAGCTAAGGCTCAAGTTGTTGGCTGGCCACCAGTTCGCTCTTACCGCAAGAACGTTATGGTTTCTTG
CCAAAAGTCTTCTGGTGGTCCAGAAGCTGCTGCTTTCGTTAAGGGAGCTGGAGCAGGAGCCGGAGCTCCAAAAGATCCAGCAAAACCAC
CAGCAAAAGCCCAGGTTGTGGGTTGGCCACCAGTGCGTTCATATCGTAAGAACGTGATGGTGTCATGTCAGAAATCATCAGGTGGTCCAG
AAGCCGCAGCCTTCGTGAAAGAGAACCTCTACTTCCAATCCGGAAAGGACTACAAGGACCACGACGGAGATTACAAGGATCACGATATC
GATTACAAGGACGACGACGACAAG

#### >3xAID amino acid sequence

GSGGGGPKDPAKPPAKAQVVGWPPVRSYRKNVMVSCQKSSGGPEAAAFVKGAGAGAKEPKDPAKPPAKAQVVGWPPVRSYRKNVMVSCQK
SSGGPEAAAFVKGAGAGAGAPKDPAKPPAKAQVVGWPPVRSYRKNVMVSCQKSSGGPEAAAFVKENLYFQSGKDYKDHDGDYKDHDIDYKD
DDDK

***Legend:***

**AID tag**
**Linker**
**TEV site**
**3x FLAG**

#### >nTIR1(*eft-3p*::TIR1::SL2::NLS::mTagBFP2::*eft-3* 3'UTR) nucleotide sequence

caacttccattggttcttccattgtttctgttaaattaatgaatttttcataaaataaagacattatacaatataaaaatgaagaatttattgaaaataaactgccagagagaaaaagtatgc
aacactcccgccgagagtgtttgaaatggtgtacggtacattttcgtgctaggagttagatgtgcaggcagcaacgagagggggagagattttttgggccttgtgaaattaacgtgagtttt
ctggtcatctgactaatcatgttggtttttgttggtttattttgtttttatctttgtttttatccagattaggaaatttaaattttatgaatttataatgaggtcaaacattcagtcccagcgtttttcctg
ttctcactgtttagtcgaatttttattttaggctttcaacaaatgttctaactgtcttatttgtgacctcactttttatattttttaatttttaaaaatattagaagtttctaggataatttttttcgacttttt
attctctctaccgtccgcactcttcttacttttaaattaaattgttttttttcagttgggaaacactttgctcactccgtagcagccATGCAAAAGCGTATAGCACTTTCATTCCC
TGAAGAGGTCCTGGAACATGTATTTTCATTCATACAGCTTGATAAGgtacgactacctgcctgcctaccgcctAAATTTTTTgtgaagTTTcttcAAAAAAtcca
gAAAAAAAAAcaaTTTTcatacgaTTTTTTcccttAAAAttgtgaaTTTTcatgcTTTTTagccccAAAAgtcattaTTTgagAAAAAATTTcatacAAAAAAgTTTTgag
AAAtacacaaTTTTTTAAAtgtaaTTTTcAAATTTTcaaTTTTcaactagAAAAttcacAAAActtgtAAATTTTggaccAAAAcaTTatacaattacTTTTTTtgaatcta
ataactacaataactacAAATTTTgttcagGATCGAAATTCAGTTTCCTTGGTTTGCAAATCATGGTATGAGATTGAGCGTTGGTGTCGTAGAAAGGTC
TTTATCGGTAATTGTTATGCTGTGTCACCAGCAACCGTAATCAGAAGATTTCCAAAGGTCAGATCAGTCGAGCTCAAGGGAAAGCCGCACTT
TGCCGACTTTAACTTGGTCCCTGATGGCTGGGGCGGATATGTATATCCATGGATCGAAGCAATGTCGTCCTCTTACACCTGGCTGGAAGAG
ATTAGATTGAAACGTATGGTTGTCACGGACGATTGCTTAGAGTTGATAGCCAAGTCTTTTAAGAATTTCAAGGTTCTTGTATTGTCCAGTTGCGA
AGGATTTTCGACGGACGGTCTTGCCGCTATTGCCGCCACATGCCGTAATCTTAAGGAATTAGACTTGCGAGAGTCCGACGTGGACGAAGT
AAGTGGACACTGGTTATCCCATTTCCCAGACACCTACACCTCGCTCGTCTCGCTCAATATATCTTGTCTCGCGAGTGAGGTGTCCTTCTCAG
CTCTCGAAAGACTCGTGACGAGATGCCCTAACCTCAAGTCTTTGAAGCTGAACCGAGCAGTGCCTCTTGAAAAATTGGCGACCCTCTTGCA
GCGTGCACCTCAGCTTGAAGAATTAGGAACCGGTGGCTACACAGCGGAGGTTCGACCTGACGTGTATAGTGGCCTTTCTGTTGCGCTTAG
TGGATGCAAAGAACTGAGATGCCTCTCTGGTTTCTGGGATGCCGTACCAGCATATCTGCCGGCCGTTTACTCTGTTTGCAGTAGACTGACC
ACACTGAACCTCTCTTACGCAACCGTGCAGTCATACGATTTGGTCAAACTGTTATGCCAATGCCCAAAACTCCAGAGACTTTGGGTTTTAGA
TTACATAGAGgtaagatatgggAAAgaaggAAAAAAccgagaTTTTacttgAAAAAttgaaTTTTTcgcgggaTTTTcaccAAAAAttgttgaatattcattaTTTcacgctg
tAAAAcTTTTAAAAAAAAaatcAAAAActacgttgAAAtcgcgTTTTTaagcgaaTTTTcttcagaattgccagaTTTTaaccccAAATTTTgcagTTTTTAAAtAAAATT
TcaccTTTTcggctcAAAttgtagaTTTTcctgAAAATTTagtacAAAAAAcaaTTTcctcgtAAATTTTTcAAATTTTcagGATGCCGGATTAGAAGTTCTGGCG
AGTACGTGCAAAGACCTTCGAGAGCTTCGTGTTTTTCCTTCCGAACCATTCGTTATGGAACCGAATGTGGCACTGACCGAGCAAGGACTGG
TCTCCGTTTCCATGGGATGCCCTAAATTAGAGTCGGTACTGTATTTCTGCCGACAAATGACTAATGCAGCCCTCATTACCATTGCAAGAAATC
GACCGAATATGACACGATTCCGATTGTGTATCATCGAGCCGAAGGCCCCGGATTACCTCACTCTGGAACCACTCGACATCGGTTTCGGTG
CGATCGTGGAGCATTGTAAGGATCTTCGTAGACTTTCATTATCCGGTTTGTTGACGGACAAAGTTTTCGAGTACATAGGAACATACGCCAAAA
AAATGGAGATGCTTTCTGTGGCTTTTGCGGGAGACAGTGATTTGGGCTTGCATCACGTGTTGAGTGGCTGTGACAGTCTGCGTAAACTGGAG
ATACGAGATTGCCCATTCGGTGACAAAGCTCTTCTTGCGAACGCTTCAAAACTCGAGACCATGCGTTCCCTCTGGATGTCTTCGTGCAGTG
TGTCCTTTGGAGCATGTAAGCTCCTTGGTCAGAAGATGCCAAAACTCAACGTGGAGGTCATTGATGAACGTGGCGCTCCAGACTCACGTC

### Supplementary Tables

CTGAGTCATGCCCGGTTGAAAGAGTCTTTATCTACAGAACCGTTGCGGGACCAAGATTCGATATGCCTGGCTTCGTTTGGAATATGGACCA
GGATTCTACTATGCGTTTTTCACGTCAGATCATCACGACGAATGGTCTCTAAgctgtctcatcctactttcacctagttaactgcttgtcttaaaatctatgcttctct
ttagtatctaaaattttcctagaagcttacaagtatataaatggtctcttctcaataaaggttgtatatttattcatcttattgaatctgccatttcctcgtttttgcgagtttatataccttcc aattt
tctttctattgtattttcaacttctaattttaattcagggaaactgcttcaacgcatcATGCCAGCTGCCAAGAGAGTCAAACTTGACATGGTATCCAAAGGAGAGG
AACTGATAAAAGAGAATATGCACATGAAGTTATATATGGAAGgtaagTTTatctAAAAAgTTTTTcattcAAAAtgtgtAAAAAttcaTTTAAAAtaacccAAA
AAtcattaatcctcgataTTTTcagGAACGGTAGATAATCACCATTTCAAATGTACCTCTGAGGGTGAAGGAAAGCCATATGAAGGCACTCAGACGA
TGAGAATCAAGGTCGTAGAAGGTGGACCTCTGCCATTCGCCTTCGATATACTTGCCACCTCCTTCTTGTATGGTTCAAAAACCTTTATAAACC
ACACGCAGGGTATTCCTGATTTTTTCAAGCAGTCTTTTCCTGAGGGATTCACATGGGAACGAGTAACTACATACGAGGACGGAGGTGTCCTG
ACAGCAACACAAGATACTTCATTACAAGATGGTTGTCTTATATATAATGTGAAGATAAGAGgtaTTTTcctgcaTTTTTcaactgggAAAAtgAAAgAAA
AtcgataaTTTcagGTGTCAATTTCACGAGTAACGGCCCGGTGATGCAGAAAAAGACGTTAGGCTGGGAGGCGTTCACAGAGACGCTTTATCC
AGCTGATGGTGGACTCGAGGGCAGAAATGATATGGCTCTCAAGTTGGTTGGAGGATCGCATCTTATTGCAAATGCCAAAACAACATATAGAT
CTAAAAAACCGGCCAAAAACCTTAAGATGCCTGGCGTCTACTACGTGGATTATCGTCTCGAACGAATCAAAGAAGCTAATAACGAGACTTAT
GTCGAGCAACACGAGGTAGCTGTAGCGCGATATTGCGATTTGCCATCAAAGCTTGGACATAAACTTAATTAAatcttcattgttgagtttatcttgttgattt
ttgaataaattatcaactctttactttttaatgggttatgaaataaataaacattgaaaactgataaacaacgttcatctct

***Legend:***

**eft-3 promoter**
**nTIR1**
**SL2 (gpd-2/3 intergenic sequence)**
**c-Myc Nuclear Localization Signal (NLS)**
**mTagBFP2**
**eft-3 3'UTR**

**>nTIR1(*eft-3p*::TIR1::SL2::NLS::mTagBFP2::*eft-3* 3'UTR) amino acid sequence**

MQKRIALSFPEEVLEHVFSFIQLDKDRNSVSLVCKSWYEIERWCRRKVFIGNCYAVSPATVIRRFPKVRSVELKGKPHFADFNLVPDGWGGYVYP
WIEAMSSSYTWLEEIRLKRMVVTDDCLELIAKSFKNFKVLVLSSCEGFSTDGLAAIAATCRNLKELDLRESDVDEVSGHWLSHFPDTYTSLVSLNIS
CLASEVSFSALERLVTRCPNLKSLKLNRAVPLEKLATLLQRAPQLEELGTGGYTAEVRPDVYSGLSVALSGCKELRCLSGFWDAVPAYLPAVYSVC
SRLTTLNLSYATVQSYDLVKLLCQCPKLQRLWVLDYIEDAGLEVLASTCKDLRELRVFPSEPFVMEPNVALTEQGLVSVSMGCPKLESVLYFCRQM
TNAALITIARNRPNMTRFRLCIIEPKAPDYLTLEPLDIGFGAIVEHCKDLRRLSLSGLLTDKVFEYIGTYAKKMEMLSVAFAGDSDLGLHHVLSGCDSL
RKLEIRDCPFGDKALLANASKLETMRSLWMSSCSVSFGACKLLGQKMPKLNVEVIDERGAPDSRPESCPVERVFIYRTVAGPRFDMPGFVWNM
DQDSTMRFSRQIITTNGL*MPAAKRVKLDMVSKGEELIKENMHMKLYMEGTVDNHHFKCTSEGEGKPYEGTQTMRIKVVEGGPLPFAFDILATSFL
YGSKTFINHTQGIPDFFKQSFPEGFWERVTTYEDGGVLTATQDTSLQDGCLIYNVKIRGVNFTSNGPVMQKKTLGWEAFTETLYPADGGLEGRND
MALKLVGGSHLIANAKTTYRSKKPAKNLKMPGVYYVDYRLERIKEANNETYVEQHEVAVARYCDLPSKLGHKLN*

***Legend:***

**nTIR1**
**c-Myc Nuclear Localization Signal (NLS)**
**mTagBFP2**
